# Upgraded CRISPR/Cas9 Tools for Tissue-Specific Mutagenesis in *Drosophila*

**DOI:** 10.1101/2020.07.02.185652

**Authors:** Gabriel T. Koreman, Qinan Hu, Yineng Xu, Zijing Zhang, Sarah E. Allen, Mariana F. Wolfner, Bei Wang, Chun Han

## Abstract

CRISPR/Cas9 has emerged as a powerful technology for tissue-specific mutagenesis. However, tissue-specific CRISPR/Cas9 tools currently available in *Drosophila* remain deficient in three significant ways. First, many existing gRNAs are inefficient, such that further improvements of gRNA expression constructs are needed for more efficient and predictable mutagenesis in both somatic and germline tissues. Second, it has been difficult to label mutant cells in target tissues with current methods. Lastly, application of tissue-specific mutagenesis at present often relies on Gal4-driven Cas9, which hampers the flexibility and effectiveness of the system. Here we tackle these deficiencies by building upon our previous CRISPR-mediated tissue restricted mutagenesis (CRISPR-TRiM) tools. First, we significantly improved gRNA efficiency in somatic tissues by optimizing multiplexed gRNA design. Similarly, we also designed efficient dual-gRNA vectors for the germline. Second, we developed methods to positively and negatively label mutant cells in tissue-specific mutagenesis by incorporating co-CRISPR reporters into gRNA expression vectors. Lastly, we generated genetic reagents for convenient conversion of existing Gal4 drivers into tissue-specific Cas9 lines based on homology-assisted CRISPR knock-in (HACK). In this way, we expand the choices of Cas9 for CRISPR-TRiM analysis to broader tissues and developmental stages. Overall, our upgraded CRISPR/Cas9 tools make tissue-specific mutagenesis more versatile, reliable, and effective in *Drosophila*. These improvements may be also applied to other model systems.

## INTRODUCTION

The ability to characterize gene function in a tissue-specific manner has been critical for studying developmental and disease mechanisms of essential genes. The clustered regularly interspaced short palindromic repeats (CRISPR)/Cas9 system has recently provided powerful tools for inducing tissue-specific gene loss of function (LOF). In this system, the endonuclease Cas9 is directed by a small guide RNA (gRNA) to a specific DNA sequence to create double-strand breaks (DSBs) (1). In the absence of homologous repair templates, DSBs are primarily repaired by non-homologous end joining (NHEJ), an error-prone process that often introduces mutations in the form of insertions or deletions (indels) (2, 3). Because the protospacer adjacent motif (PAM) required for Cas9 action is ubiquitous in genomes (1, 4), by targeting the expression of Cas9 and gRNAs to specific tissues, mutations can be induced at virtually any gene in a tissue-specific manner. However, current tissue-specific CRISPR/Cas9 tools in *Drosophila* are still deficient in three areas, limiting the power of CRISPR/Cas9 in analyzing gene functions in broad tissues and biological processes.

### Method of tissue-specific Cas9 delivery

In *Drosophila*, CRISPR/Cas9-mediated tissue-specific mutagenesis is generally achieved by two approaches that differ in the method of Cas9 delivery. The first approach uses a tissue-specific Gal4 to drive *UAS-Cas9* expression, and expresses gRNAs using either a ubiquitous or a UAS promoter (5, 6). The vast number of available tissue-specific Gal4 lines (1, 7–9) makes adoption of this method relatively easy. For this reason, Gal4-driven Cas9s have been successfully used to elucidate gene functions, such as in circadian rhythm (10, 11), and to screen for new genes involved in neuronal remodeling (12).

The second method, CRISPR-mediated tissue-restricted mutagenesis (CRISPR-TRiM), relies on enhancer-driven Cas9 for tissue specificity, and employs ubiquitously expressed gRNAs (13). Compared to the *Gal4/UAS-Cas9* approach, CRISPR-TRiM has several advantages. First, enhancer-driven Cas9 involves only one transcription step and thus requires less time for expression than Gal4-driven Cas9, reducing the chance of perduring gene products masking defects of mutant cells (13). Therefore, CRISPR-TRiM is more effective for studying early phenotypes of mutant cells. Second, enhancer-driven Cas9 is usually expressed at much lower levels than Gal4-driven Cas9, alleviating cytotoxicity associated with high Cas9 expression (13). Third, CRISPR-TRiM is a simpler system that requires only two genetic components, facilitating the construction of tissue-specific knockout strains with fewer time-consuming crosses. Lastly, CRISPR-TRiM allows simultaneous use of Gal4/UAS for manipulating other tissues. This flexibility of CRISPR-TRiM was demonstrated by simultaneous neuronal gene knockout (KO) and Gal4-dependent labeling of phosphatidylserine exposure in neurodegeneration (14). Until now, CRISPR-TRiM has been limited by the small number of tissue-specific Cas9 lines currently available. Wide applications of CRISPR-TRiM in *Drosophila* require efficient ways of generating new Cas9 lines that are specific to various tissues and developmental stages.

### gRNA efficiency

Successful tissue-specific mutagenesis requires efficient transgenic gRNAs, as inefficient gRNAs would result in uneven LOF in the target tissue and complicate the analysis. A sound general strategy for improving gRNA efficiency is to optimize the design of gRNA expression vectors. So far, optimizations have been made mainly in two areas. First, since expressing multiple gRNAs targeting a single gene can increase the likelihood of mutagenesis (5, 15), considerable efforts have been devoted to making multi-gRNA (multiplexed) expression vectors (5, 6, 16). Studies in rice, *Drosophila*, and yeast have demonstrated the effectiveness of tRNA-gRNA designs for the efficient expression and processing of multiplexed gRNAs (5, 17, 18). In these designs, multiple gRNAs are interspaced by glycine (G) tRNAs (tRNA^Gly^) in a single transcript under the control of a single promoter. Endogenous tRNA-processing enzymes cut out tRNAs from the transcript, simultaneously releasing individual gRNAs.

A second aspect of gRNA optimization concerns the scaffold sequence that forms hairpin loops to complex with Cas9 (1, 19). In an early study, a modified scaffold containing a flip of A-U positions and a stem-loop extension (F+E) was found to improve the targeting of Cas9 to the intended locus (20). More recently, an additional extension of the second stem-loop (gRNA2.1) was found to further increase the mutagenic efficiency of gRNAs in human cells (21). To develop general strategies for making highly efficient gRNAs for tissue-specific mutagenesis in *Drosophila*, we previously combined tRNA^Gly^-gRNA with the (F+E) gRNA scaffold in a transgenic gRNA vector. This vector performed much more efficiently than previous gRNA vector designs in somatic tissues (13). However, there is still room to further improve the design of gRNA vectors towards higher gRNA efficiency and more reliable tissue-specific mutagenesis. Moreover, the germline differs from the soma in important ways that often impact transgene expression (22, 23). It has thus been unknown which design is the most efficient in the *Drosophila* germline for use in germline mutagenesis and gene replacement through homology-directed repair (HDR).

### Labeling of mutagenized cells

An unsolved caveat of all current methods of CRISPR-mediated mutagenesis is the inability to label mutant cells in the target tissue. This is particularly problematic for data analysis when Cas9 activity is not evenly distributed across all cells in the tissue of interest. This problem cannot be solved simply by fusing Cas9 to a fluorescent protein because the presence or absence of Cas9 protein at the time of analysis does not necessarily correlate with the presence or absence of mutations. Therefore, the ability to label mutant cells in the target tissue is an unmet need.

Here, we present our strategies to tackle these challenges. We report further improvements in the design of gRNA vectors that lead to higher gRNA efficiency and more reliable tissue-specific mutagenesis in somatic tissues. We also address germline performance of various constructs and report the most efficient vector for germline mutagenesis. Moreover, to label mutant cells, we developed a co-CRISPR reporter system and demonstrate its applications in mutagenizing the *Drosophila* epidermis in conjunction with positive- and negative-labeling. Lastly, we generated genetic tools for convenient conversion of existing Gal4 lines into tissue-specific Cas9 lines. These news tools significantly increase the power of tissue-specific mutagenesis in *Drosophila* and make more reliable and more sophisticated CRISPR/Cas9 manipulations available for the study of broader biological questions.

## RESULTS

### A new multi-gRNA design greatly improves the efficiency of somatic mutagenesis

We previously identified an efficient multi-gRNA design (tgFE) that employs both tRNA^Gly^ as spacers to separate gRNAs and the (F+E) gRNA scaffold to enhance gRNA/Cas9 interaction (13). We have continued to optimize this design to further increase its mutagenic efficiency. Reasoning that tRNA processing could affect the rate and level of gRNA production, we first tested alternative tRNAs in conjunction with the (F+E) gRNA scaffold. We compared six *Drosophila* tRNAs in a dual-gRNA design, in which a *Drosophila* tRNA^Gly^ is followed by an irrelevant gRNA (targeting the blue fluorescent protein BFP) and a second variable tRNA is followed by a gRNA targeting *Syntaxin 5* (*Syx5*) (Figure 1C). We previously found in an RNAi screen that *Syx5* is required for dendrite growth of *Drosophila* class IV dendritic arborization (C4da) neurons (unpublished). Therefore, the most efficient gRNA construct should yield the most robust and consistent dendrite reduction (Figures 1A and 1B). For comparison, we used a tgFE-based dual-gRNA construct that expresses two gRNAs against *Syx5* (designated as GG). When combined with a C4da-specific Cas9 (*ppk-Cas9*), only the construct containing glutamine (Q) tRNA (tRNA^Gln^) consistently caused strong (59%) reduction of dendrite length, while other versions caused much more variable reductions as indicated by the deviation of each sample from the mean dendrite length (Figures 1C and 1D). As *Syx5* is required for ER to Golgi transport (24) and is likely expressed early in the neuronal lineage, we speculate that the variability of dendrite reduction is due to variable timings of mutagenesis in post-mitotic neurons. If so, incorporating tRNA^Gln^ may have led to faster processing of multiplexed gRNAs and therefore a more consistent depletion of Syx5 protein.

**Figure 1:**
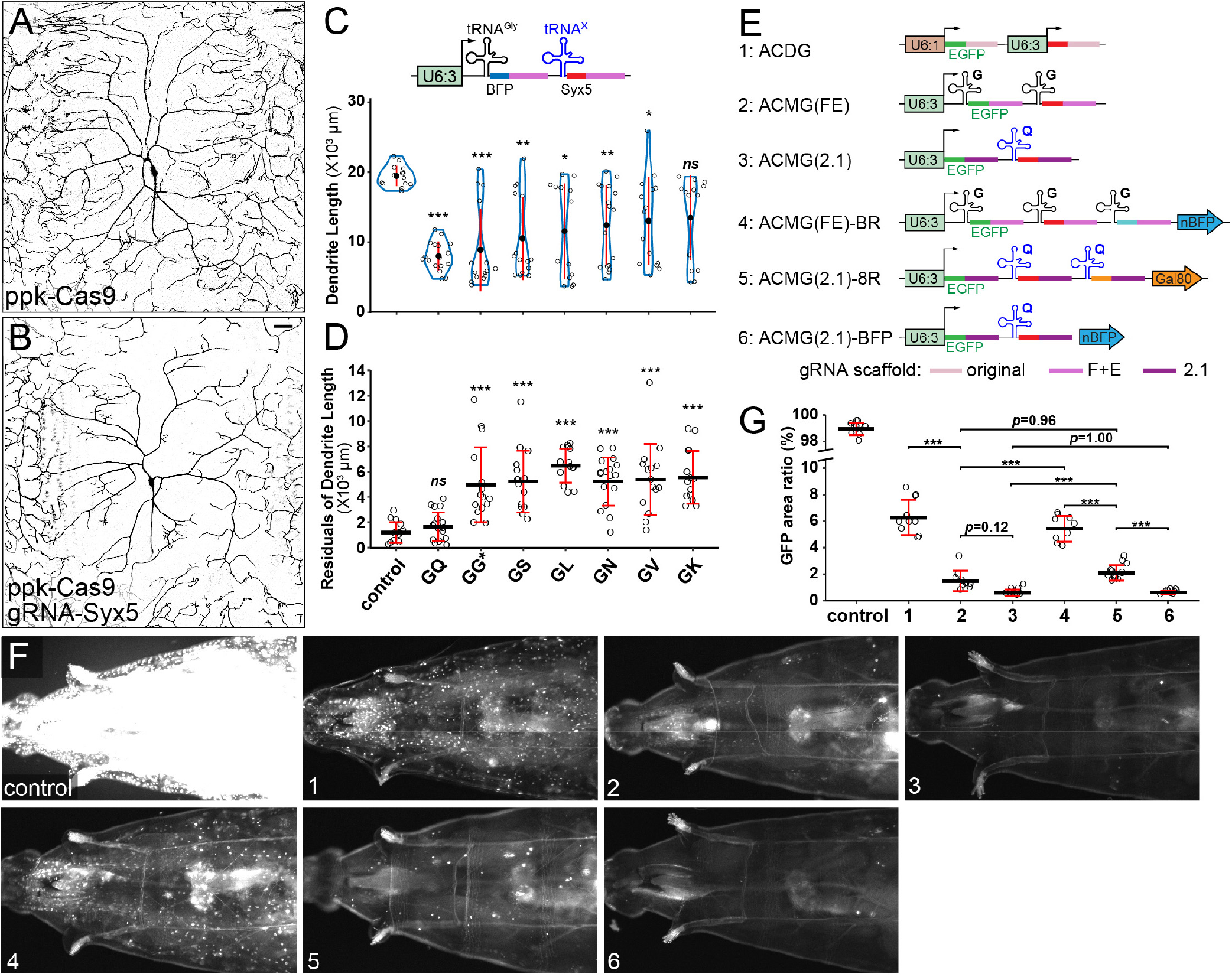
A new multi-gRNA design significantly improves the efficiency of somatic mutagenesis. (A and B) C4da neurons in *ppk-Cas9* control (A) and *ppk-Cas9 gRNA-BFP-Syx5(GQ)* (B). Scale bars, 50 um. (C and D) Total dendrite lengths (C) and residuals (differences from the mean) of dendrite lengths (D) of C4da neurons using the various *Syx5* gRNAs. Asterisk indicates that both gRNAs in GG target *Syx5*. The significance level above each column indicates comparison with the control. ***p ≤ 0.001, **p ≤ 0.01, *p ≤ 0.05, ns, not significant; Welch’s ANOVA and Welch’s t-tests with Bonferroni correction (C) and one-way ANOVA and Tukey’s HSD test (D). Each circle represents one neuron. n = number of neurons: GG (n = 15), GK (n = 14), GL (n = 14), GN (n = 17), GQ (n = 17), GS (n = 15), GV (n = 15), Control (n = 14). (E) Six variants of gRNA constructs targeting EGFP/GFP. Each construct contains one gRNA targeting EGFP and one targeting GFP. The identities of tRNAs are indicated and the gRNA scaffolds are color-coded. (F) EGFP patterns in larvae expressing a ubiquitous EGFP marker, *Act5C-Cas9*, and a *gRNA-EGFP* whose identity is indicated by the number defined in (E). The control larva has no gRNA construct. (G) Ratios of EGFP(+) areas in larvae. ***p ≤ 0.001, Welch’s ANOVA and Welch’s t-tests with Bonferroni correction. Numbering corresponds to gRNAs in (E). n=number of larvae: Control (n = 10); 1 (n = 10); 2 (n = 9); 3 (n = 10); 4 (n = 9); 5 (n = 12); 6 (n = 9). Each circle represents one larva. For all quantifications, Black bar, mean; red bars, SD.

We further compared tRNA variants in a quadruple-gRNA design to knock out *Nsf1* and *Nsf2* simultaneously (Figure S1A-S1C); these genes act redundantly to permit dendrite growth of C4da neurons (13). However, using tRNA^Gln^ in various combinations did not significantly enhance dendrite reduction (Figure S1D), possibly because the limiting factor in *Nsf1/Nsf2* neuronal KO is the timing of Cas9 expression rather than tRNA processing (13).

We next asked whether incorporating the gRNA2.1 scaffold could augment gRNA mutagenic efficiency relative to the (F+E) scaffold. We generated a dual-gRNA construct against EGFP/GFP, with tRNA^Gln^ as the spacer and gRNA2.1 as the scaffold (Figure 1E, ACMG(2.1)), and compared it to two earlier versions of gRNA-EGFP/GFP (Figure 1E, ACDG and ACMG(FE)). All three versions express two gRNAs targeting EGFP and GFP coding sequences separately. ACDG uses two U6 promoters and the original scaffold while ACMG(FE) is based on the tgFE design. Using a ubiquitous nuclear EGFP and *Act5C-Cas9*, we assayed the efficiency of gRNAs in knocking out EGFP in larvae. Consistent with our previous comparisons in da neurons (13), ACMG(FE) is significantly more efficient than ACDG: While ACDG removed EGFP from most cells, there were still many EGFP-positive nuclei (6.28% area) (Figures 1F and 1G). ACMG(FE) further reduced EGFP signals to only some muscle stripes and the larval brain (1.49% area) (Figures 1F and 1G). In comparison, ACMG(2.1) almost completely eliminated EGFP, leaving only occasional EGFP-positive cells (0.60% area) (Figures 1F and 1G). These data suggest that tRNA^Gln^-gRNA2.1 (referred to as Qtg2.1) is a superior gRNA design over previous generations.

It was unclear whether increasing the number of gRNAs in the multiplex design would impact the efficiency of individual gRNAs. To answer this question, we added an extra irrelevant gRNA, along with a ubiquitous reporter (described later), to ACMG(FE) and ACMG(2.1) constructs (Figure 1E, ACMG(FE)-BR and ACMG(2.1)-8R, respectively). These two triple-gRNA constructs performed worse than their dual-gRNA counterparts, although the new Qtg2.1 design was still consistently better than the tgFE version (with 2.10% and 5.42% areas, respectively) (Figures 1F and 1G). The reduced efficiencies of triple-gRNA constructs were not due to the ubiquitous reporters, as adding only a reporter to ACMG(2.1) did not change the mutagenic efficiency (0.61% area) (Figures 1F and 1G). These data suggest that adding more gRNAs could reduce the efficiency of each gRNA in a multi-gRNA design, perhaps due to competition for Cas9.

We also tested the Qtg2.1 design in quadruple/quintuple gRNA constructs against *Nsf1/Nsf2*, but neither the Qtg2.1 design nor the addition of an irrelevant gRNA significantly affected the level of dendrite reduction in C4da neurons (Figure S1D). These results are consistent with the idea that Cas9 expression timing, rather than the rate of gRNA production, is the limiting factor in this particular case (13).

In summary, we found that the Qtg2.1 gRNA design that incorporates tRNA^Gln^ and the gRNA2.1 scaffold significantly enhances mutagenic efficiency of multiplexed gRNAs.

### Co-CRISPR visualizes mutant cells in epithelial tissues

The inability to label mutant cells has limited the analytic power of current methods of tissue-specific CRISPR mutagenesis. A possible solution is to incorporate a ubiquitous reporter, as well as a gRNA targeting the reporter, into the gRNA vector (Figure 2A). In such a co-CRISPR design, LOF of the gene of interest (GOI) could be correlated with the loss of reporter expression. As a proof of concept, we tested an EGFP/GFP gRNA construct that also carries a BFP gRNA and a ubiquitous nuclear BFP (nBFP) reporter (ACMG(FE)-BR in Figure 1E). This construct theoretically should allow negative labeling of *EGFP* mutant cells by the absence of BFP signal. Pairing this construct with ubiquitous nuclear EGFP, we examined the correlation of EGFP and BFP KO in larval epidermal cells (Figure 2C). We measured the ratio of EGFP and BFP double-positive cells in all EGFP-positive epidermal cells (overlap/EGFP ratio), which should have a value of 1 when mutagenesis of the two genes is correlated exactly. Deviations from this optimal ratio would be caused by false positives, where cells reporting CRISPR/Cas9 activity (the lack of BFP) still express functional EGFP.

**Figure 2.**
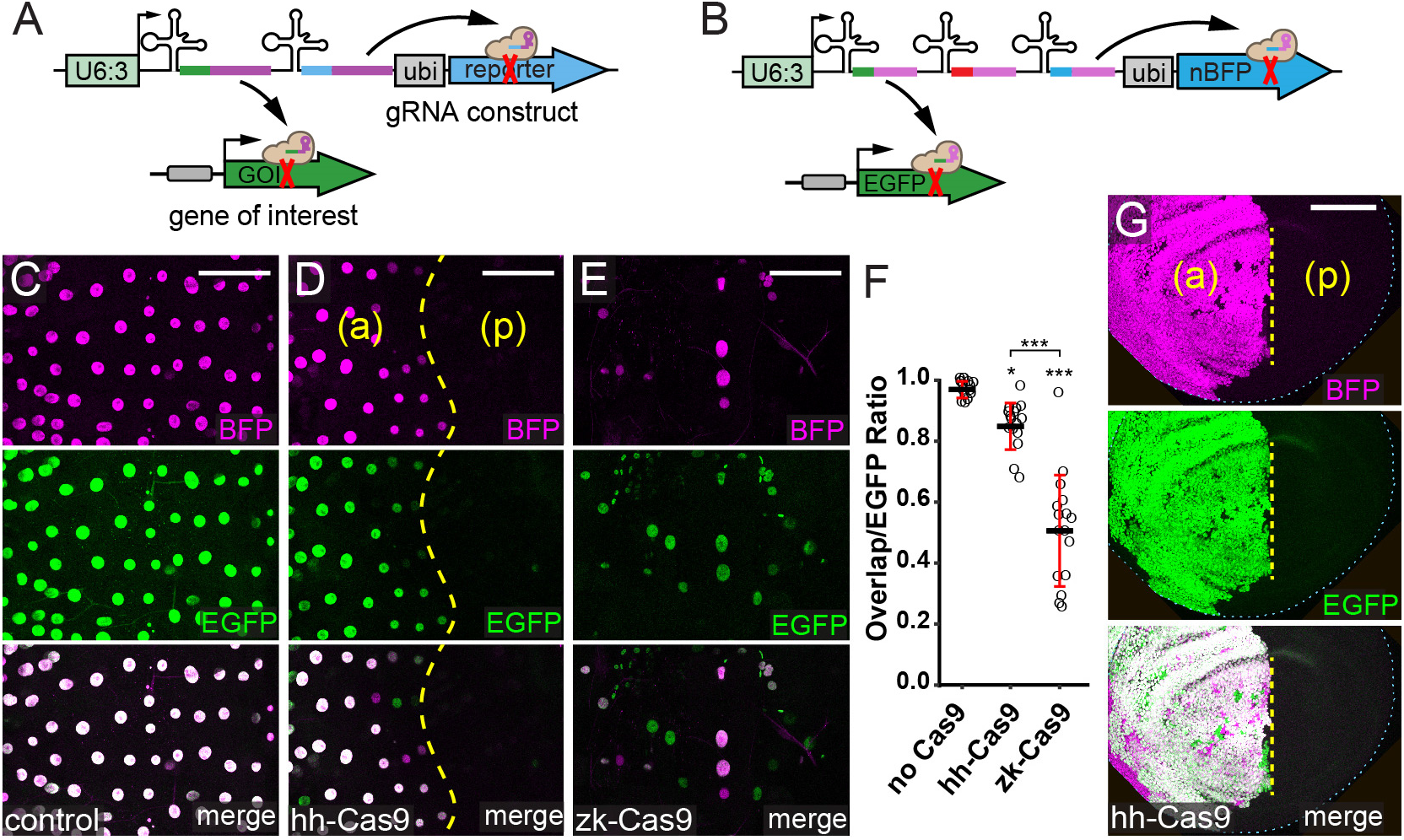
Co-CRISPR visualizes mutant cells in epithelial tissues. (A) General strategy of labeling mutant cells for a gene of interest (GOI) using a ubiquitous co-CRISPR reporter. (B) Design of negative-labeling via co-CRISPR mutagenesis of EGFP and a nuclear BFP (nBFP) reporter (*gRNA-EGFP[ACMG(FE)-BR]*). (C-E) Epidermal expression patterns of BFP (magenta) and EGFP (green) in larvae expressing a ubiquitous EGFP marker, *gRNA-EGFP[ACMG(FE)-BR]*, and no Cas9 (C), *hh-Cas9* (D), or *zk-Cas9* (E). The anterior (a) and posterior (p) hemi-segments are indicated in (D). Scale Bar, 100 um. (F) Ratio of cells expressing both EGFP and BFP over all cells expressing EGFP in larvae carrying the various Cas9 lines. The significance level above each column indicates comparison with the control. ***p ≤ 0.001, *p ≤ 0.05, contrasts of estimated marginal means (EMMs) based on a generalized linear mixed-effects model with a binomial response. Each circle represents one segment. n = number of segments: Control (n = 16), *hh-Cas9* (n = 16), *zk-Cas9* (n = 16); biological replicates = 4 larvae per genotype. Black bar, mean; red bars, SD. (G) BFP (magenta) and GFP (green) expression pattern in a wing disc of a larva expressing ubiquitous EGFP, *gRNA-EGFP[ACMG(FE)-BR]*, and *hh-Cas9*. The anterior (a) and posterior (p) compartments are indicated. Scale Bar, 100 um.

We compared two Cas9 lines, *hh-Cas9* and *zk-Cas9*, which have distinct spatiotemporal expression patterns in the larval epidermis, in co-CRISPR labeling of *EGFP* KO cells. *hh-Cas9* is expected to be expressed in the posterior half of each segment from early embryogenesis to late larval stages (13, 25). While EGFP and BFP were completely knocked out in posterior hemi-segments by *hh-Cas9*, some anterior cells also lost either EGFP or BFP (Figure 2D), amounting to an overall overlap/EGFP ratio of 0.85 (Figure 2G). We also tested *hh-Cas9* in co-CRISPR in the wing imaginal disc, as it is expressed in the posterior compartment of the wing pouch (13). Similarly, we saw complete KO of both EGFP and BFP in the posterior wing disc as expected, and sporadic KO of either EGFP or BFP in small, random patches in the anterior wing disc. The *hh-Cas9* activity in the anterior epidermal hemi-segment and the anterior wing disc is likely due to leaky expression during development, which should be transient and low. In comparison, the *zen-kr* enhancer in *zk-Cas9* is expected to drive transient expression in precursor cells of epidermal cells during early embryogenesis (26–28). It knocked out EGFP and BFP in some, but not all, epidermal cells (Figure 2E), generating an overlap/EGFP ratio of 0.51 (Figure 2F). These results suggest that reliable co-CRISPR requires (moderately) high and persistent expression of Cas9 in the cell lineage, such as *hh-Cas9* in the posterior epidermal hemi-segment and in the posterior compartment of the wing disc.

### Co-CRISPR enables positive and negative labeling of mutant cells in dendrite development and epithelial morphogenesis

To test if co-CRISPR can be used to visualize cells carrying biallelic mutations of endogenous genes, we first designed a positive-labeling gRNA construct to target *sulfateless* (*sfl*), which encodes a heparan sulfate-glucosamine N-sulfotransferase required by epidermal cells to support local growth of C4da dendrites (28). Besides expressing two gRNAs targeting *sfl*, this gRNA construct also carries a gRNA against *Gal80* and a ubiquitously expressed Gal80 reporter (Figure 3A). As Gal80 suppresses Gal4 transcription factor activity (29), the loss of *Gal80* can be visualized by Gal4-driven expression of a fluorescent marker, thus enabling positive labeling of *sfl* mutant cells. We paired this construct with *hh-Cas9*, *UAS-tdTom*, and a pan-epidermal Gal4 driver. Accordingly, we observed that tdTom-positive epidermal cells in the posterior hemi-segment lacked coverage by high order dendritic branches of C4da neurons (Figure 3B), a phenotype associated with epidermal *sfl* LOF (28). Although some tdTom-positive cells were also observed in the anterior hemi-segment next to the segment boundary, reflecting leaky expression of *hh-Cas9*, high order branches of C4da neurons were also absent from these tdTom patches. The reliability of positive labeling of *sfl* mutant cells is demonstrated by a tight correlation (Pearson’s correlation coefficient *r* = 0.96) between labeled areas with dendrite reduction and all labeled areas (Figure 3C). Overall, 95%±4% (n=29) labeled areas showed strong dendrite reduction, demonstrating that positive labeling by co-CRISPR is an effective approach for visualizing mutant epidermal cells.

**Figure 3:**
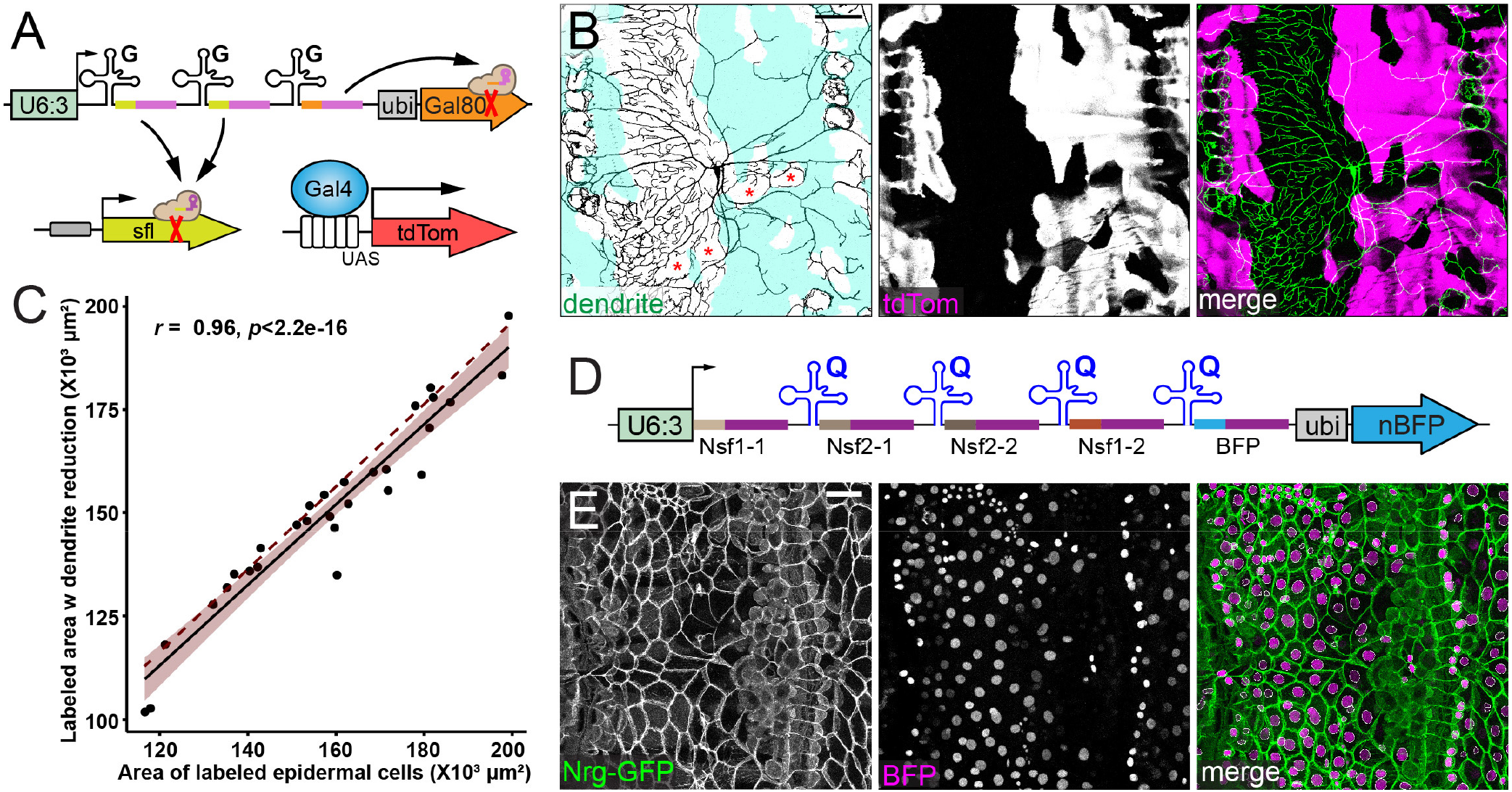
Co-CRISPR enables positive and negative labeling of mutant cells in dendrite development and epithelial morphogenesis. (A) Design of positive-labeling of *sfl* mutant cells using co-CRISPR mutagenesis of a ubiquitous Gal80 co-CRISPR reporter (*gRNA-sfl(8R)*). The labeling of Gal80-mutant cells is enabled by Gal4-driven *UAS-tdTom* expression. (B) Dendrite morphology and tdTom expression in a larva expressing *ppk-CD4-tdGFP* (green), *A58>tdTom* (magenta), *gRNA-sfl(8R)*, and *hh-Cas9*. tdTom-labeled areas are shaded in the dendrite panel. Red asterisks indicate muscle attachment cells, which C4da neurons do not innervate. Scale Bar, 100 um. (C) Correlation between tdTom-labeled areas showing dendrite reduction and all tdTom-labeled areas. The solid line shows the linear regression, the shaded area is 95% confidence interval. The dotted line indicates a perfect correction (slope = 1). Pearson’s Correlation Coefficient r = 0.96, p < 2.2 × 10^−16^. Each dot represents a segment; n = 29, biological replicates = 15. (D) Design of negative labeling of *Nsf1*/*Nsf2* double mutant cells by a nBFP co-CRISPR reporter (*gRNA-NSF1-NSF2[QQQ(2.1)-BR]*). (E) A representative image of epidermal cell morphology in larvae expressing *gRNA-NSF1-NSF2[QQQ(2.1)-BR]*, *hh-Cas9*, and *Nrg-GFP* (Green). Scale Bar, 100 um. n = number of segments = 5, biological replicates = 3.

We next further validated the effectiveness of negative labeling with the nBFP reporter by targeting *Nsf1/Nsf2* (Figure 3D). Epidermal cells lacking both *Nsf1* and *Nsf2* delaminate from the epidermal sheet and become round (13). Because this phenotype requires successful KO of all four alleles in the cell, reliable labeling of double mutant cells is expected to be more challenging. To increase the mutagenic efficiency, we used the Qtg2.1 design in this construct, as opposed to the tgFE design in *EGFP/GFP* and *sfl* co-CRISPR reporter constructs. When paired with *hh-Cas9*, this construct generated many BFP-negative cells in the posterior epidermal hemi-segment, all of which showed deformed morphology such as delamination and round shape (Figure 3E).

Together, our results suggest that both positive- and negative-labeling co-CRISPR constructs reliably report mutagenic events in the larval epidermis when paired with a proper Cas9.

### gRNA mutagenic efficiency is governed by different rules in the germline and the soma of *Drosophila*

As the tgFE design has a superior mutagenic efficiency in somatic tissues than multi-gRNAs driven by separate Pol III promoters (13), we wondered if this principle is also true in the *Drosophila* germline. Therefore, we compared multiple versions of gRNA constructs to knock out three genes known to play important roles in germline development and/or early embryogenesis of the progeny. *bam* is required for germline development in both males and females and its loss blocks oogenesis in females and suppresses spermatogenesis in males (15, 30). *cid* is required for centromere identity in meiosis (31) and its LOF reduces male fertility (15). The third gene, *gnu*, is maternally required for early embryogenesis (32); thus, eggs derived from the germline of *gnu* mutant females show reduced hatchability (33). For each gene, we designed two constructs based on the tgFE design, with one containing a potent Pol III promoter CR7T (15) and the other containing the U63 promoter (#1 and #2 in Figure 4A, respectively). As a comparison, for each gene, we also generated a dual-gRNA construct driven by separate CR7T and U6:3 promoters (CR7T-U63(FE), #3 in Figure 4A).

**Figure 4:**
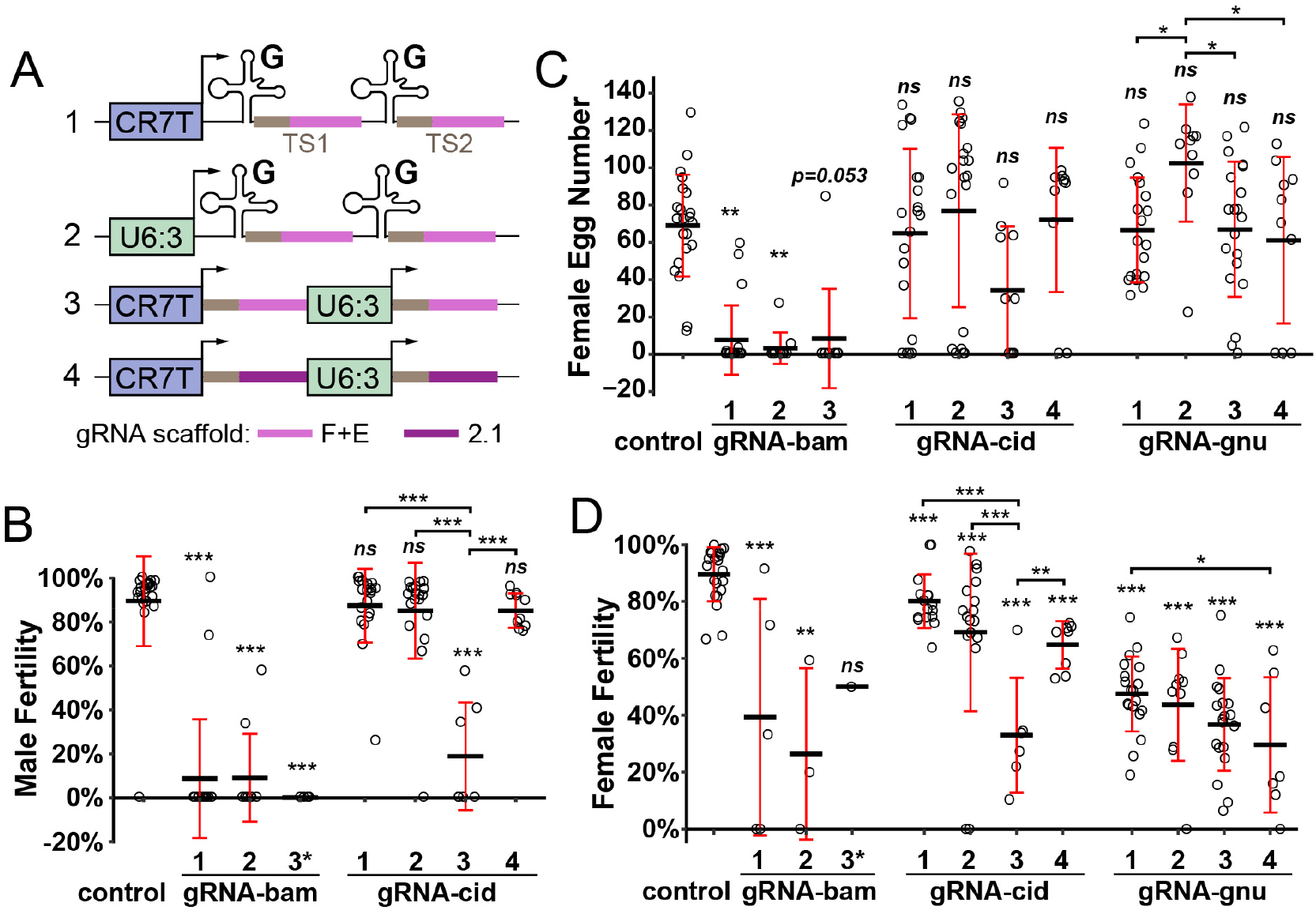
gRNA mutagenic efficiency is governed by different rules in the germline and the soma of *Drosophila*. (A) Design of gRNA constructs tested in the germline. Numbering of gRNAs corresponds to that in (B-D). (B) Hatchability of eggs derived from crosses using males expressing gRNAs targeting *bam* or *cid*. Control had no gRNAs. Contrasts of estimated marginal means (EMMs) based on proportion of hatched/unhatched eggs using a generalized linear mixed-effects model with a binomial response. The asterisk on *gRNA-bam(3)* indicates that Fisher’s Exact Test was used to compare this group with the control as it had no variance. n = number of single-male × single-female crosses. Control (n = 22); bam-1 (n = 20); bam-2 (n = 10); bam-3 (n = 5); cid-1 (n = 20); cid-2 (n = 20); cid-3 (n = 7); cid-4 (n = 10). (C) Number of eggs laid by females expressing gRNAs targeting *bam*, *cid*, or *gnu*. Control had no gRNAs. Contrasts of estimated marginal means (EMMs) using a generalized linear model with a negative binomial response. n = number of single-male x single-female crosses. Control (n = 22); bam-1 (n = 20); bam-2 (n = 10); bam-3 (n = 10); cid-1 (n = 20); cid-2 (n = 20); cid-3 (n = 10); cid-4 (n = 10); gnu-1 (n = 19); gnu-2 (n = 10); gnu-3 (n = 20); gnu-4 (n = 10). (D) Hatchability of eggs derived from females expressing gRNAs targeting *bam*, *cid*, or *gnu*. Control had no gRNAs. Contrasts of estimated marginal means (EMMs) based on proportion of hatched/unhatched eggs using a generalized linear mixed-effects model with a binomial response. The asterisk indicates that *gRNA-bam(3)* is not significantly different from the control due to its small sample size, as only one female laid eggs. n = number of single-male x single-female crosses. Control (n = 22); bam-1 (n = 5); bam-2 (n = 3); bam-3 (n = 1); cid-1 (n = 16); cid-2 (n = 17); cid-3 (n = 6); cid-4 (n = 8); gnu-1 (n = 19); gnu-2 (n = 10); gnu-3 (n = 19); gnu-4 (n = 7). For all quantifications, the significance level above each column indicates comparison with the control. **p ≤ 0.01, *p≤ 0.05, ***p ≤ 0.001, ns, not significant; Black bar, mean; red bars, SD.

We carried out KO experiments using a germline-specific Cas9, *nos-Cas9* (6), and measured “male fertility”, which is the percentage of hatched eggs from individual wildtype females mated with the KO males, “female egg number”, which is the number of eggs laid by individual KO females, and “female fertility”, which is the percentage of hatched eggs from individual KO females mated with wildtype males. We only measured female egg number and female fertility for the *gnu* KO, as *gnu* does not play a role in the male germline. All three constructs for *bam* yielded strong reduction of male fertility and female egg number, but surprisingly, with CR7T-U63(FE) showing the highest percentage of complete loss of fertility in both males and females (100% and 90%, respectively) (Figures 4B and 4C). This suggests that the efficiency of *bam* gRNAs are high and not limited by the construct design. For *cid*, we found that the CR7T-U63(FE) design was also much more efficient than tgFE designs in reducing male fertility and female fertility (Figures 4B and 4D): While CR7T-U63(FE) reduced 79% of male fertility and 63% of female fertility, the tgFE constructs did not affect male fertility and reduced 11%-23% of female fertility. For *gnu*, although differences among the three constructs were not statistically significant for female fertility, the CR7T-U63(FE) version again appeared to be the most efficient (Figure 4D). These data indicate that including tRNAs in the gRNA construct negatively impacts mutagenic efficiency in the germline.

We then wondered whether gRNA2.1 can enhance gRNA mutagenic efficiency in the germline. Since the *bam* gRNAs were already very efficient, we made another dual-promoter-dual-gRNA construct using the gRNA2.1 scaffold (CR7T-U63(2.1), #4 in Figure 4A) for *cid* and *gnu* only. While this version appeared to increase the efficiency for *gnu* slightly (Figure 4D), it is significantly worse than CR7T-U63(FE) for *cid* (Figures 4B and 4D). Although this inconsistency could be specific to the target sequences used in our constructs due to RNA secondary structures, our data nevertheless suggest that CR7T-U63(FE) is the most reliable gRNA design for germline mutagenesis. Importantly, our results show that efficient mutagenesis requires different gRNA design principles in the germline and the soma, likely due to differences in RNA processing.

### Gal4-to-Cas9 conversion expands the options for tissue-specific Cas9

Our CRISPR-TRiM strategy relies on tissue-specific Cas9 lines, the limited availability of which is currently the bottleneck for wide applications of CRISPR-TRiM. Meanwhile, a large number of tissue-specific Gal4 lines are available for virtually every tissue and all developmental stages of *Drosophila*. To take advantage of the Gal4 resources and eliminate the need for cloning and transgenesis for making new tissue-specific Cas9 lines, we developed a fast and reliable way to convert existing Gal4 lines into Cas9 lines through a few simple genetic crosses (Figure S2), based on homology assisted CRISPR knock-in (HACK) (34). This conversion utilizes a Cas9-donor transgenic construct that carries a 2A-Cas9 coding sequence flanked by Gal4 homology arms, a CR7T-U63(2.1) dual-gRNA cassette targeting the Gal4 coding sequence, and a ubiquitous nBFP marker for separating the Cas9-donor insertion from the converted Cas9 (Figure 5A). When combined with *nos-Cas9* and the Gal4 line of interest, the gRNAs generate DNA double-strand breaks in the middle of the Gal4 coding sequence and allow in-frame incorporation of 2A-Cas9 via homology-directed repair. The “self-cleaving” 2A peptide will release Cas9 and a truncated and nonfunctional Gal4 as two separate proteins after translation. The converted Cas9 chromosome can be detected by a positive Cas9 tester that we previously reported (13) (Figure S2).

**Figure 5:**
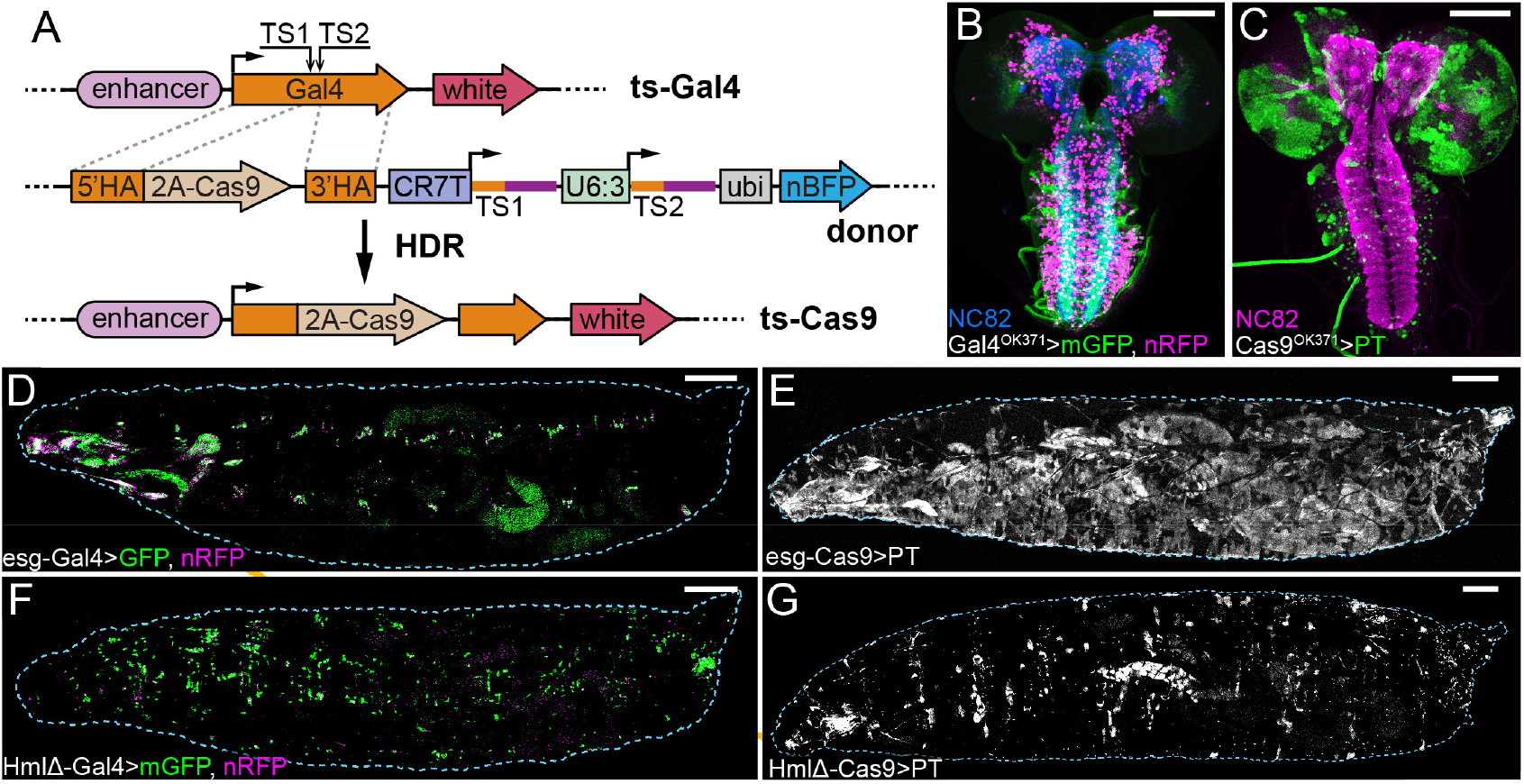
Gal4-to-Cas9 conversion expands the choices of tissue-specific Cas9. (A) Diagram of Gal4-to-Cas9 conversion using a HACK donor line. The donor expresses two gRNAs (TS1 and TS2) targeting the tissue-specific (ts) Gal4, which results in in-frame incorporation of 2A-Cas9 into the Gal4 locus through homology-directed repair (HDR). The donor expresses *ubi-nBFP* that can be selected against when screening for Cas9 convertants. (B and C) Activity patterns of *OK371-Gal4* (B) and *OK371-Cas9* (C) in the larval brain. NC82 staining shows brain neuropils. (D and E) Activity patterns of *esg-Gal4* (D) and *esg-Cas9* (E) in whole larvae. (E and F) Activity patterns of *HmlΔ-Gal4* (D) and *HmlΔ-Cas9* (E) in whole larvae. Gal4 activity patterns were visualized by UAS-driven expression of CD4-tdGFP (mGFP) or GFP (green) and nuclear RFP (nRFP, magenta). Cas9 activity patterns were visualized by Cas9 positive tester (PT). Scale Bars, 100 um in (B) and (C), 250 um in (D-G).

As a proof of principle, we converted three Gal4 lines that are specific to different tissues into Cas9 lines. We compared activity patterns of the Cas9 lines, as revealed by the Cas9 positive-tester *Act5C-Gal4 UAS-GFP; tub-Gal80 gRNA-gRNA* (13), to those of the original Gal4 lines. Interestingly, Cas9s and the corresponding Gal4s do not always show identical activity patterns. *OK371-Gal4* is specific to glutamatergic motor neurons in the third instar ventral nerve cord (VNC) (Figure 5B) (35), while *OK371-Cas9* activity was detected in only a subset of these motor neurons, as well as in sporadic glial cells in the brain lobes (Figure 5C). *esg-Gal4* is active in stem cell populations including larval histoblasts (Figure 5D) (36), but we detected a much broader activity pattern of *esg-Cas9* in the larval epidermis (Figure 5E). Lastly, while *HmlΔ-Gal4* is specific to larval hemocytes (Figure 5F) (37), the activity of *HmlΔ-Cas9* was detected in hemocytes and also a small number of random epidermal cells (Figure 5G). The discrepancies between Gal4 and Cas9 activity patterns are likely due to two factors. First, whereas Gal4 reporters only reflect the current Gal4 activity, the Cas9 positive tester reports accumulated Cas9 activity in the entire cell lineage during development, which could lead to broader Cas9 patterns. Second, labeling of Cas9 activity by the positive tester relies on complete elimination of Gal80 protein and therefore is subject to *Gal80* perdurance, which could result in more restricted labeling than the true Cas9 pattern. Despite these differences, our method of Gal4-to-Cas9 conversion simplifies the generation of new tissue-specific Cas9 lines and opens up broader opportunities for using CRISPR-TRiM to study gene function.

## DISCUSSION

CRISPR/Cas9-mediated mutagenesis holds great promise in advancing genetic analysis in *Drosophila* and is currently undergoing rapid development (38–40). However, existing tools for tissue-specific mutagenesis can still be improved to increase their power for discovery and functional analysis of genes. First, essential to the reliability of LOF analysis, gRNA efficiency can be further improved for both somatic and germline mutagenesis. Second, methods for labeling mutant cells within the tissue of interest are still missing. Lastly, although Gal4/UAS-Cas9 can be adopted for tissue-specific mutagenesis, convenient ways of applying CRISPR/Cas9 independent of Gal4/UAS are needed to maximize the simplicity, flexibility, and effectiveness of the system. In this study, we present solutions to these deficiencies. The new tools we developed will likely be useful for *Drosophila* researchers to address broad biological questions and can be adapted to improve CRISPR/Cas9 approaches in other systems.

The success of tissue-specific mutagenesis depends vitally on gRNA efficiency. Although many algorithms have been developed to predict gRNA efficiency based on the target sequence (41), optimized gRNA expression vectors are still needed for achieving the maximal gRNA efficiency. The design of the gRNA expression vector can affect the rate of gRNA production and gRNA-Cas9 complex formation. We previously found that the tgFE design that incorporates tRNA^Gly^ spacers and the (F+E) gRNA scaffold for making multiplexed gRNAs was more efficient than other designs in knocking out genes in neurons (13). Now we show that the combination of tRNA^Gln^ and the gRNA2.1 scaffold (the Qtg2.1 design) further improves gRNA efficiency in broad somatic tissues. This increase of gRNA efficiency is especially important for knocking out genes in polyploid tissues like muscles and glia or when more gRNAs are expressed simultaneously. It will also likely facilitate unmasking LOF phenotypes of genes expressed early in the cell lineage.

Surprisingly, we found that including tRNAs in multi-gRNA constructs is detrimental for germline mutagenesis. This unexpected result may reflect differences of tRNA processing mechanisms in somatic and germline tissues. In addition, although gRNA2.1 worked well in somatic tissues, our data suggest that it is comparable or worse than (F+E) in the germline. Therefore, our results demonstrate that mutagenesis in the soma and the germline requires different optimizations of the gRNA expression vector. For maximal efficiency, we recommend dual-gRNAs based on the Qtg2.1 design for somatic mutagenesis while we prefer dual-gRNAs with the (F+E) scaffold driven by separate promotors for germline mutagenesis.

To solve the challenge of labeling mutant cells in tissue-specific mutagenesis, we incorporated co-CRISPR systems in gRNA vectors to report Cas9 activity in the tissue of interest both positively and negatively. Due to the nature of CRISPR/Cas9-mediated mutagenesis, mutations in the targeted cells are inherently heterogeneous. Therefore, a reporting system that can truly reflect the nature of mutations in every cell is unlikely to be feasible. However, we show that a practical approach is to correlate the loss of the target gene with that of a co-CRISPR reporter. Because this approach requires reliable mutagenesis of both the GOI and the reporter, its success depends on both the Cas9 and the gRNAs for the GOI. First, Cas9 needs to be expressed evenly and at a relatively high level in the intended tissue so that both the GOI and the reporter have ample opportunities to be mutated. For example, the high and persistent expression of *hh-Cas9* in the posterior wing disc results in reliable labeling of mutant cells, while its low and transient leaky expression in the anterior wing disc cannot be faithfully reported. Second, as gRNAs for co-CRISPR reporters are already chosen to be highly efficient, gRNAs for the GOI also need to be adequately efficient to minimize false-positive reporting. In practice, both Cas9s and gRNA transgenes needed to be validated before use in labeled mutagenesis, such as by using our EGFP-BFP reporter line and the Cas9-LEThAL assay (13), respectively. It is important to note that it is not appropriate to predict the quantitative degree of GOI LOF by measuring expression levels of co-CRISPR reporters, as incomplete removal of co-CRISPR reporters indicates weak or late Cas9 activity and hence poor correlation between mutagenesis of GOI and the reporters. Therefore, phenotypic analysis should be conducted only in cells that show robust co-CRISPR labeling.

To overcome the limited availability of tissue-specific Cas9 lines, we adopted the HACK approach (34) and developed reagents for convenient conversion of existing Gal4 lines into Cas9 lines. This easy conversion involves no cloning or transformation steps. This method adds to the existing options of enhancer-driven Cas9 lines (13) and puts CRISPR-TRiM analysis within reach of the broader *Drosophila* community. As a proof of principle, we generated three tissue-specific Cas9s. Interestingly, converted Cas9s and the original Gal4s do not always show identical activity patterns, likely due to the difference in the ways their activities are visualized. It is worth noting that the option of Gal4-driven Cas9 may not alleviate this discrepancy because Gal4-driven Cas9 could also have leaky expression (39) and mutagenesis by Gal4-driven Cas9 suffers even more from perdurance (13). Nevertheless, our results suggest that it is important to validate the activity patterns of converted Cas9s before using them in tissue-specific mutagenesis.

Excitingly, large-scale transgenic gRNA libraries are currently in production and being made available to the *Drosophila* community (12, 39, 40). However, the mutagenic efficiencies of existing gRNAs vary significantly (39, 40). While progress has been made to improve gRNA efficiency in *Drosophila* and mammalian systems (13, 21), many optimizations have not been incorporated into these libraries. With the new designs we present here, future libraries could be developed to fit specific screening needs, for example, in the germline, in somatic tissues, or for marked mutagenic analysis.

## MATERIALS AND METHODS

**Table S1.**
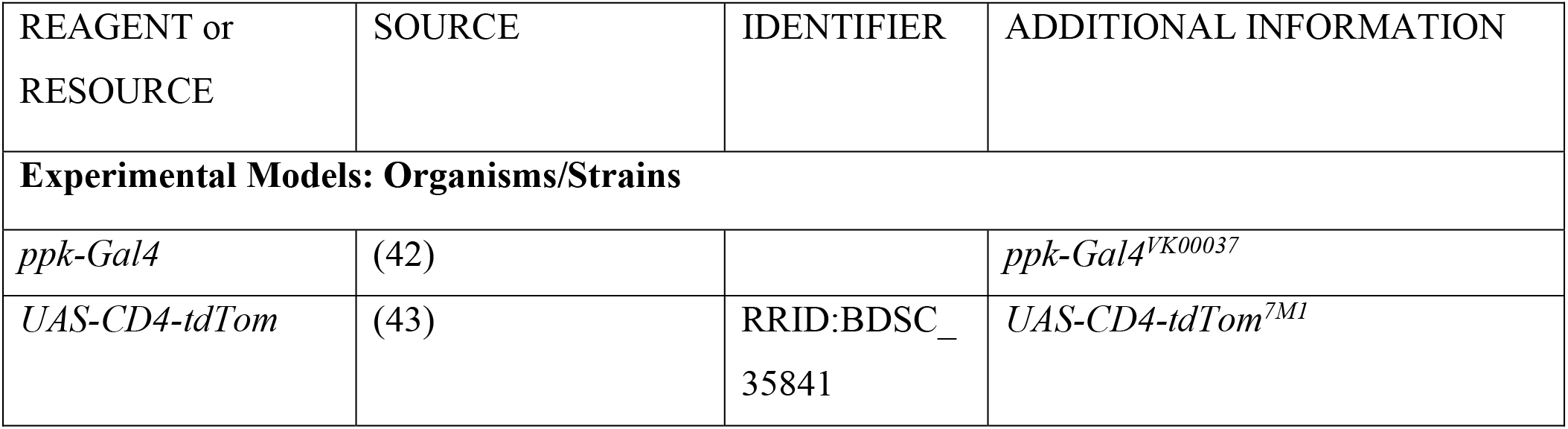
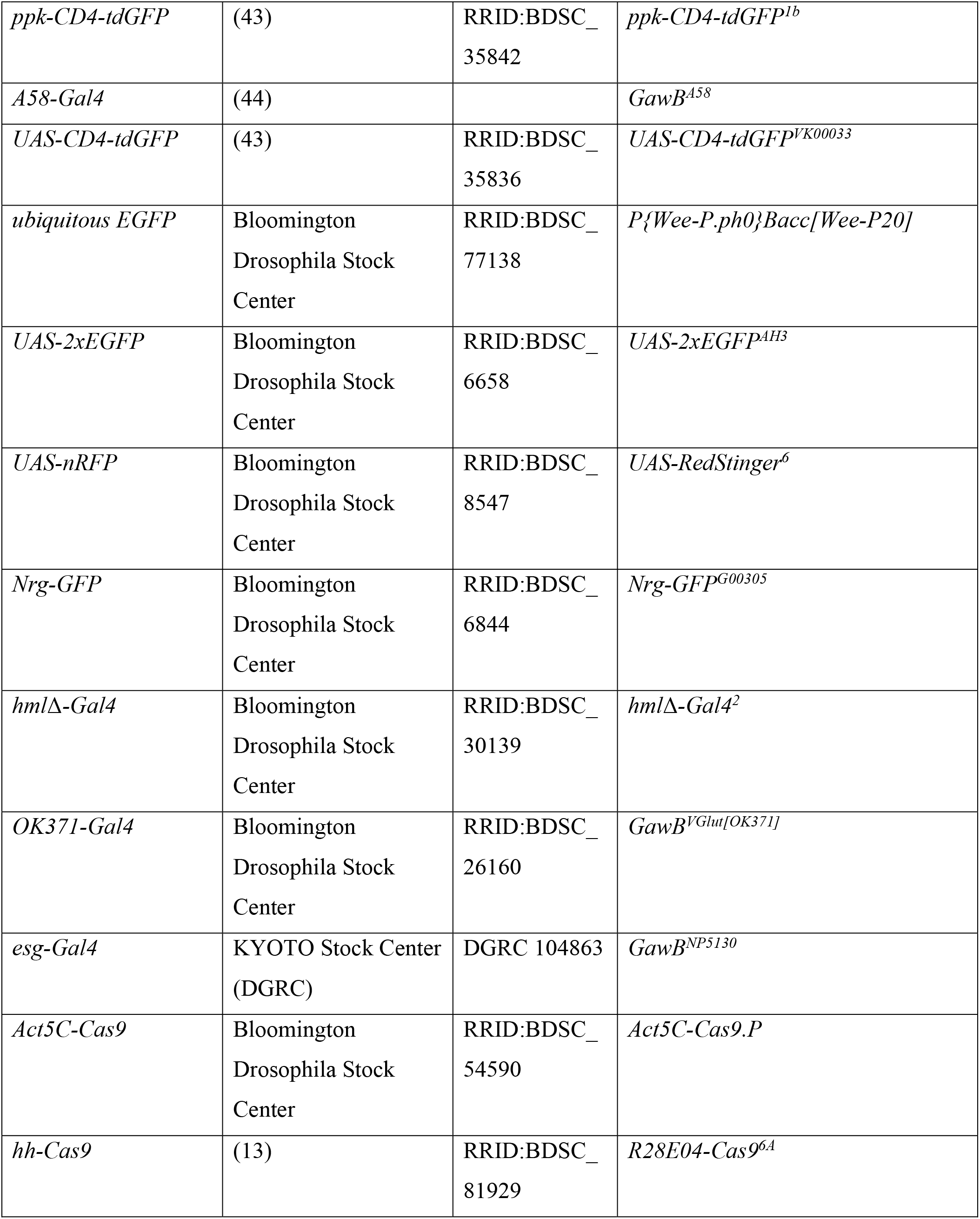
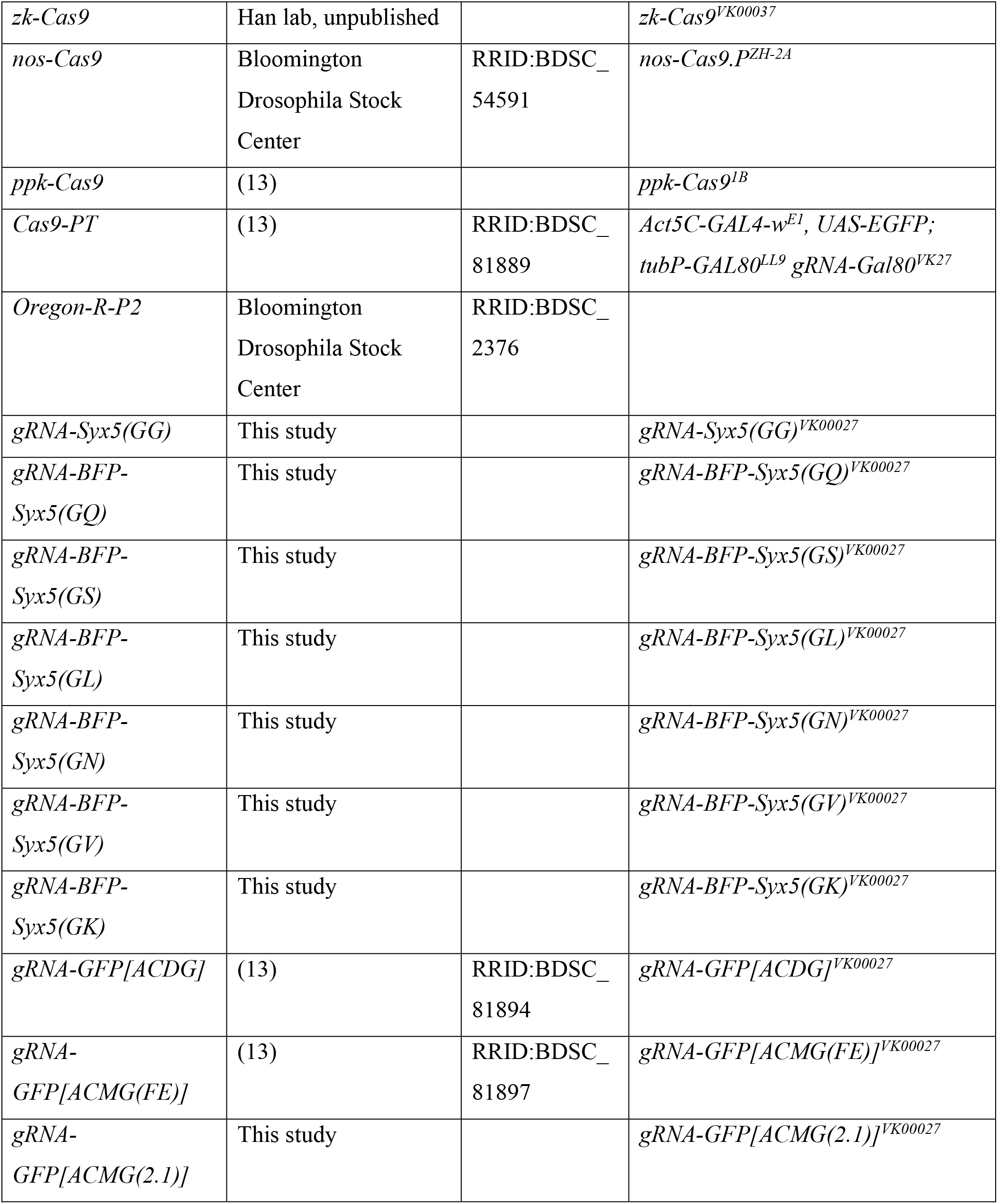
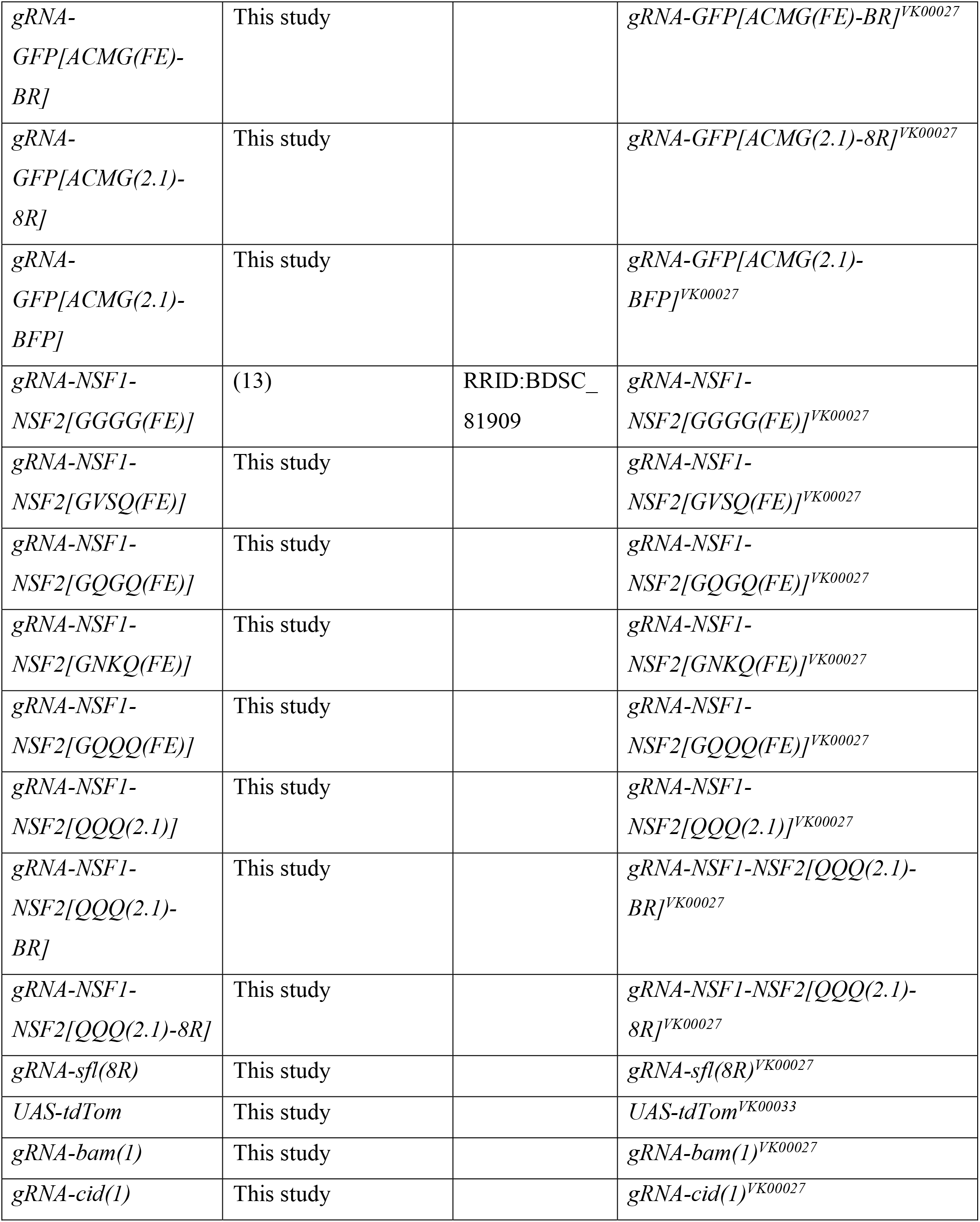
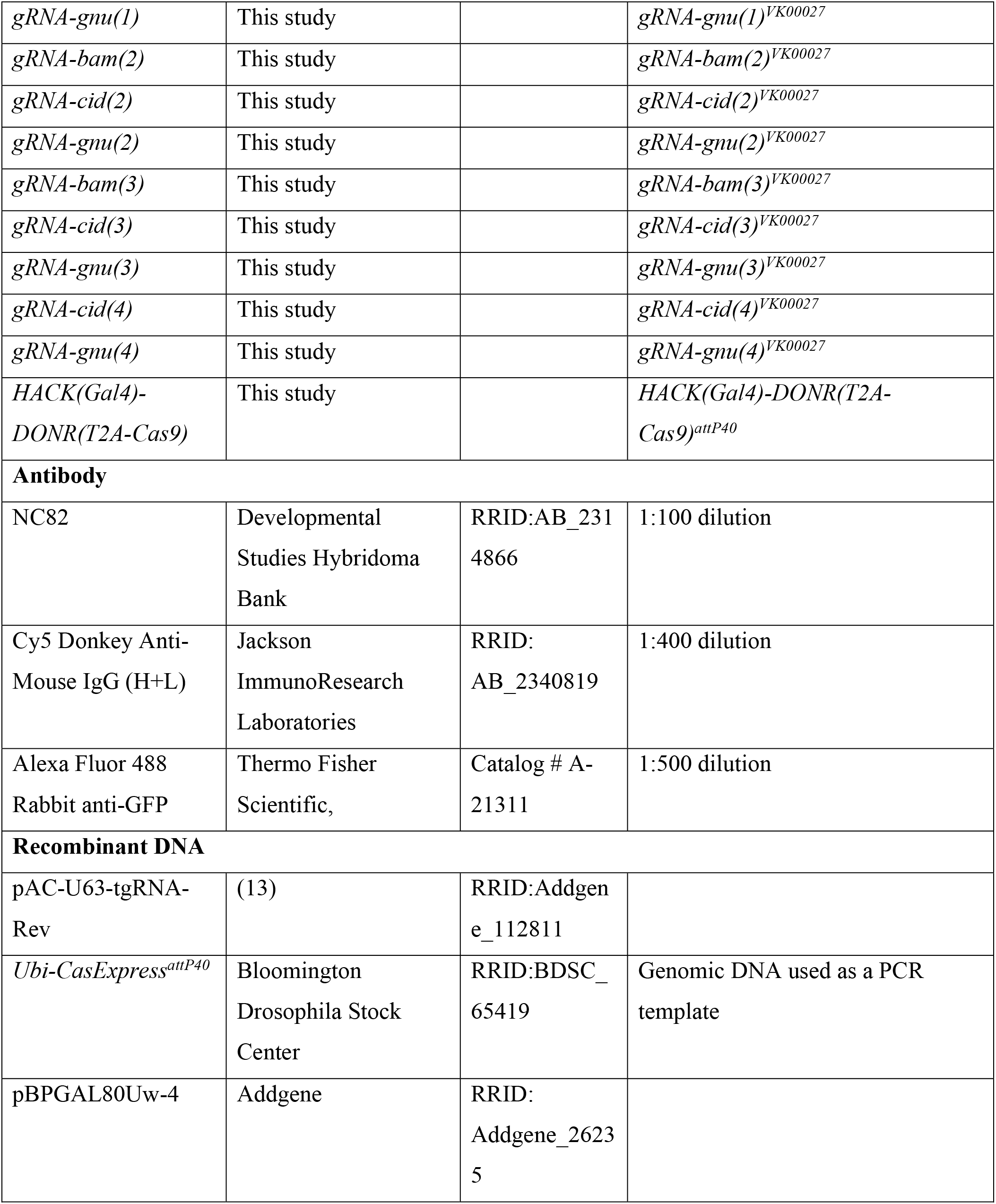
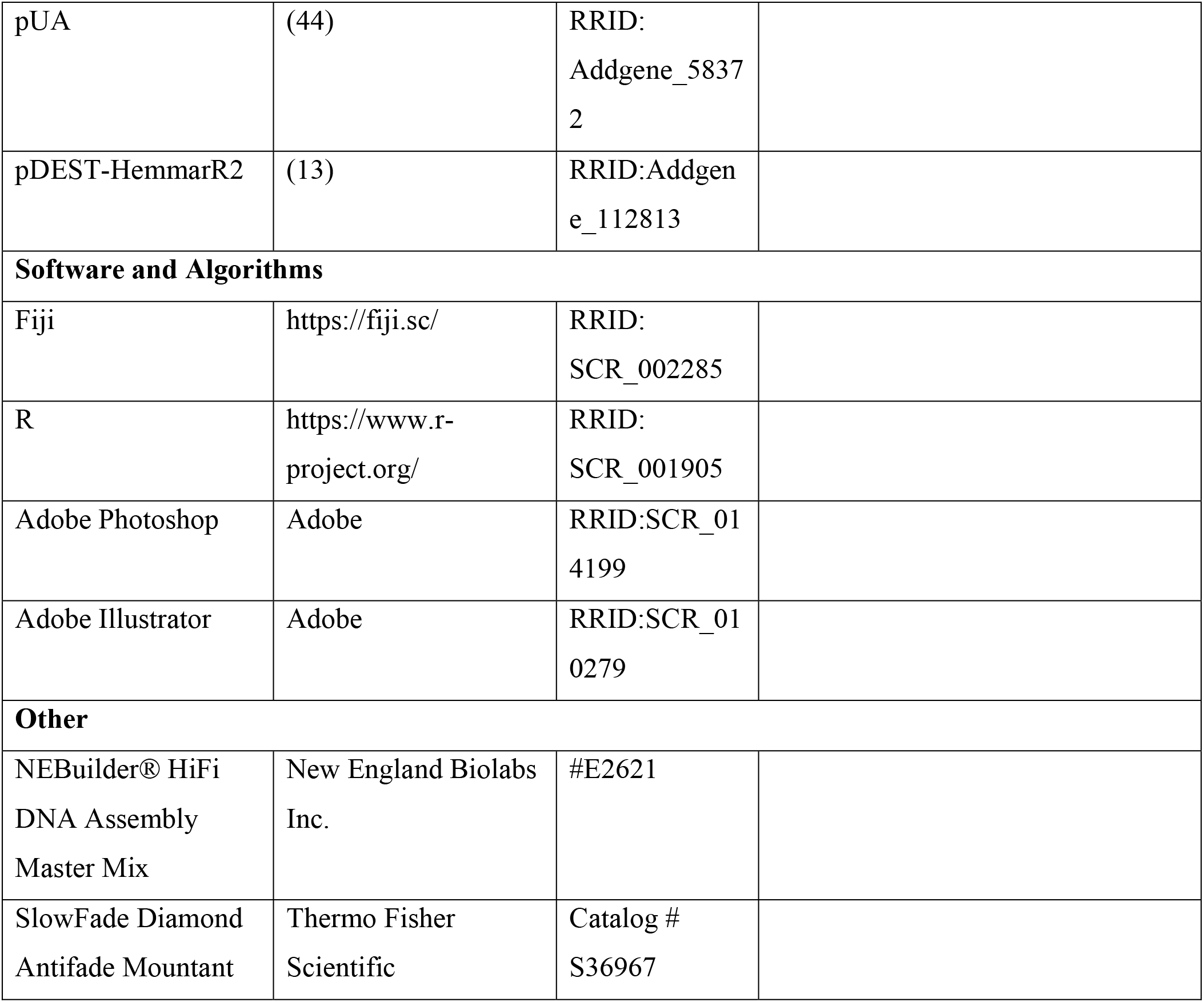
Key Resource Table.

### Fly Stocks

See Table S1 (Key Resource Table) for details of fly stocks used in this study. All flies were cultured on standard yeast-glucose medium in a 12:12 light/dark cycle at 25℃. We use *ppk-CD4-tdGFP* and *ppk>CD4-tdTom* to visualize C4da neurons; *CyO, Wee-P20* balancer as ubiquitous nuclear EGFP; *Nrg-GFP* to visualize epidermal cell shape; *UAS-CD4-tdGFP*, *UAS-RedStinger*, and *UAS-EGFP* to visualize Gal4 activity; and Cas9 positive tester (PT) to visualize Cas9 activity.

### Molecular Cloning

#### gRNA cloning vectors

Nine gRNA cloning vectors listed in Table S2 were constructed in the pAC (attB-CaSpeR4) backbone (13). Each of them contains in order some or all of the following components as specified in Table S2: a Pol III promoter, a tRNA, SapI cloning sites, a gRNA scaffold, a gRNA targeting the co-CRISPR reporter, U6 3’ flanking sequence, and a ubiquitous co-CRISPR reporter. The U6:3 promoter was PCR amplified from pAC-U63-tgRNA-Rev (Addgene #112811). The CR7T promoter was synthesized as a gBlock DNA fragment (IDT, Inc.). tRNAs and gRNA scaffolds were synthesized as gBlock DNA fragments. The promoter of *Ubi-p63E* was PCR amplified from *Ubi-CasExpress* genomic DNA (45). mTagBFP-NLS was synthesized as a gBlock DNA fragment. Gal80 coding sequence was PCR amplified from pBPGAL80Uw-4 (Addgene 26235). A *His2Av* polyA sequence after the BFP/Gal80 coding sequence was PCR amplified from pDEST-HemmarR2 (Addgene # 112813).

**Table S2.**
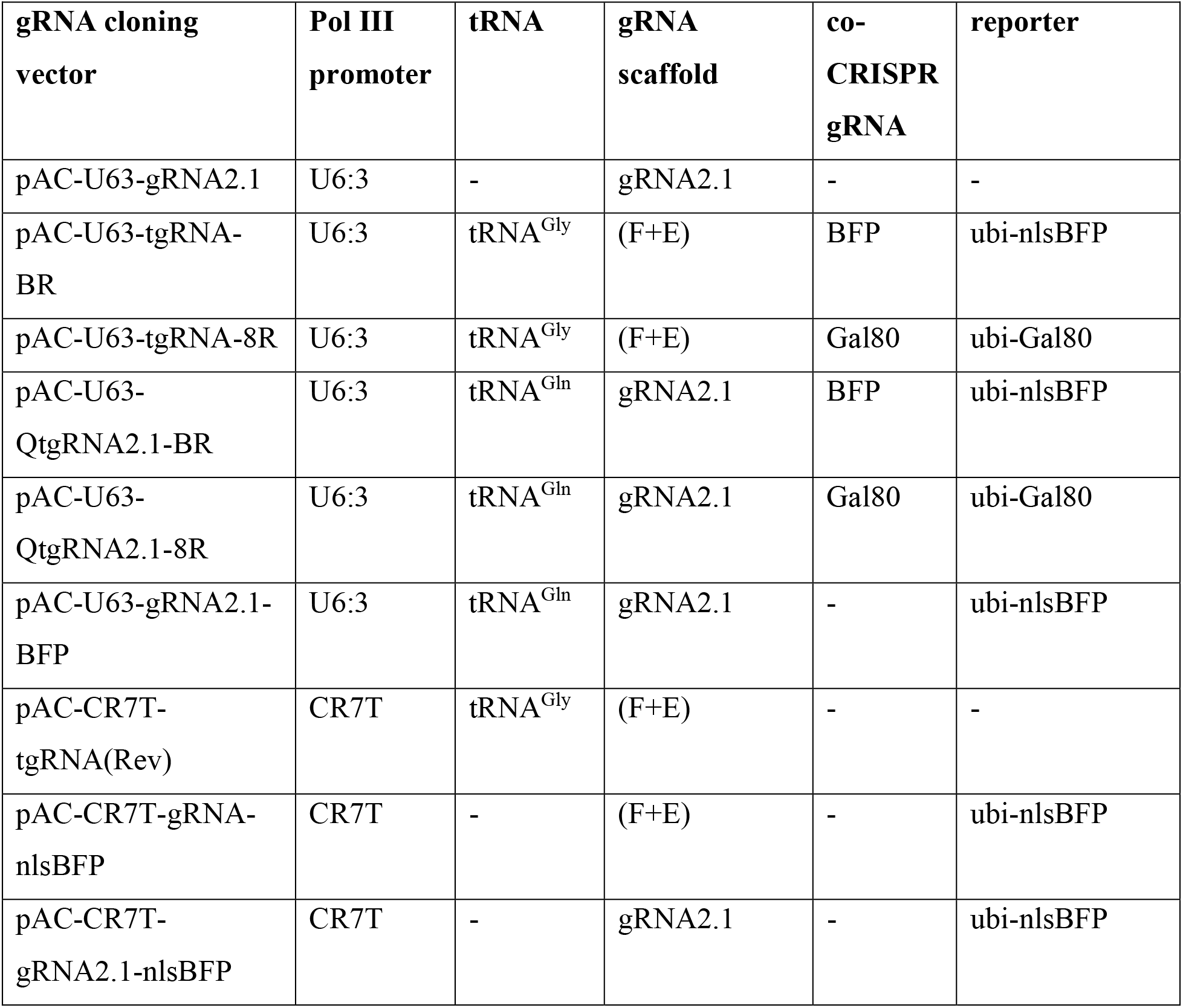
gRNA cloning vectors.

#### gRNA PCR template vectors

Eight PCR template vectors as listed in Table S3 were constructed for generating PCR fragments used for assembling the final gRNA expression vectors. Each of them was made by assembling a synthetic gBlock DNA fragment with a PCR-amplified Kanamycin resistant backbone using NEBuilder DNA Assembly (New England Biolabs). The region to be PCR-amplified in each vector contains a gRNA scaffold followed by either a tRNA or the U6:3 promoter. The sequences of tRNAs are listed in Table S4.

**Table S3.**
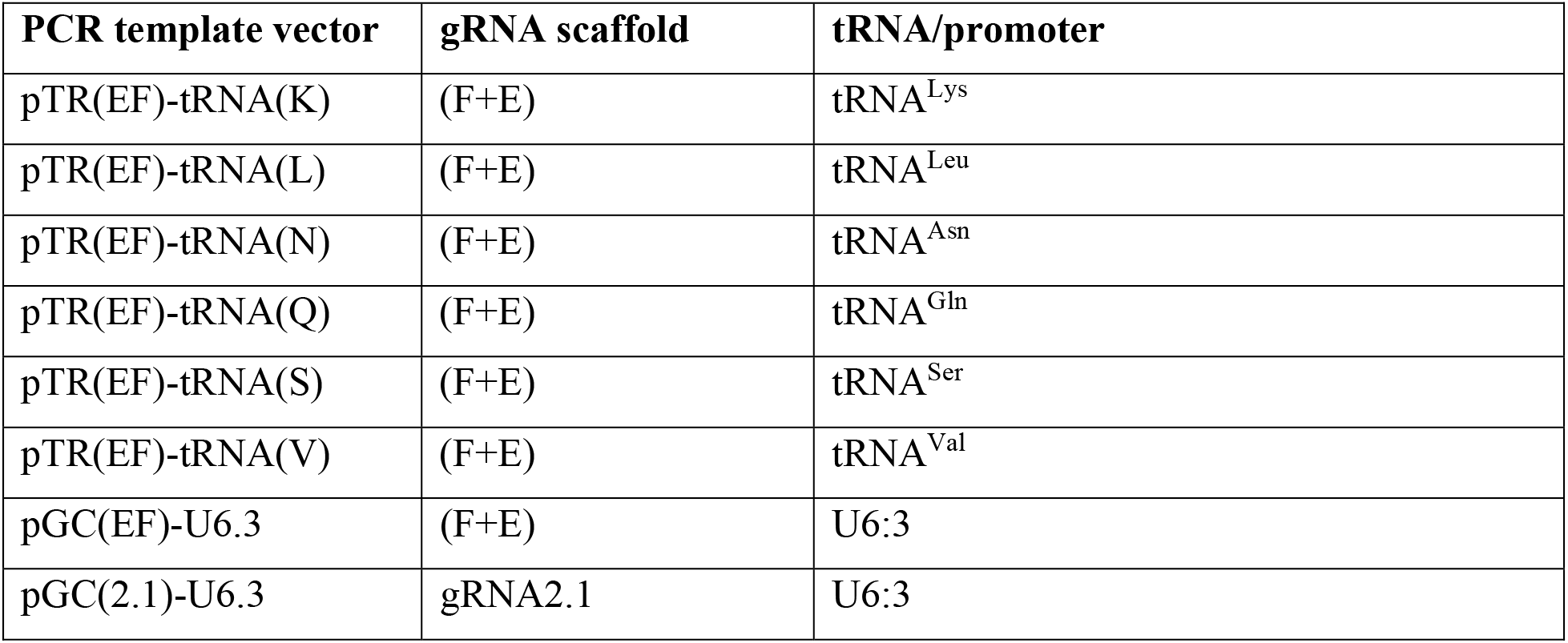
gRNA PCR template vectors.

**Table S4.**
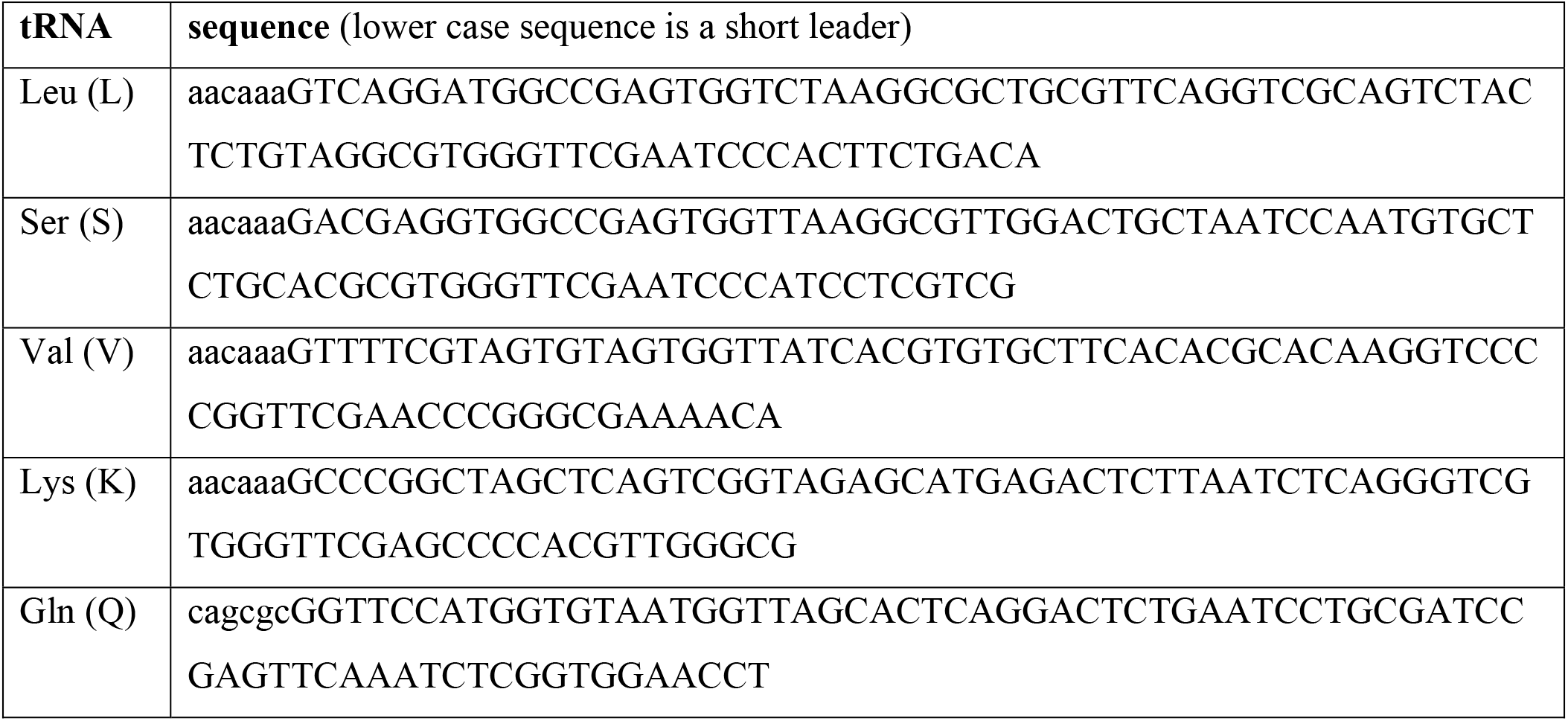
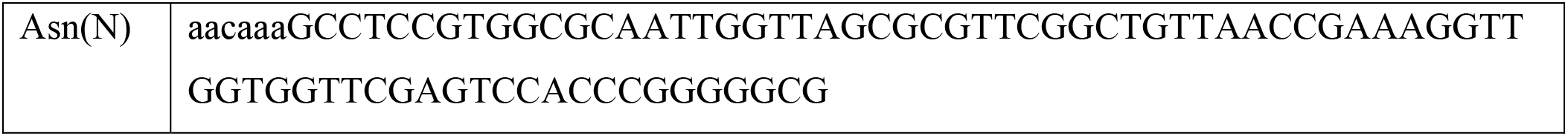
tRNA sequences.

#### gRNA expression vectors

30 gRNA expression vectors as listed in Table S5 were constructed with appropriate gRNA cloning vectors and gRNA PCR template vectors. Primers were designed to contain appropriate gRNA target sequences and sequences complementary to the PCR template vectors. The PCR products were then assembled with SapI-digested gRNA cloning vectors using NEBuilder DNA Assembly. Table S6 lists the gRNA target sequences used in this study. Table S7 provides a guideline for designing primers and choosing PCR template vectors for making Qtg2.1 multi-gRNAs and reporter gRNAs for somatic tissues and CR7T-U6:3 dual-gRNA constructs for the germline.

**Table S5.**
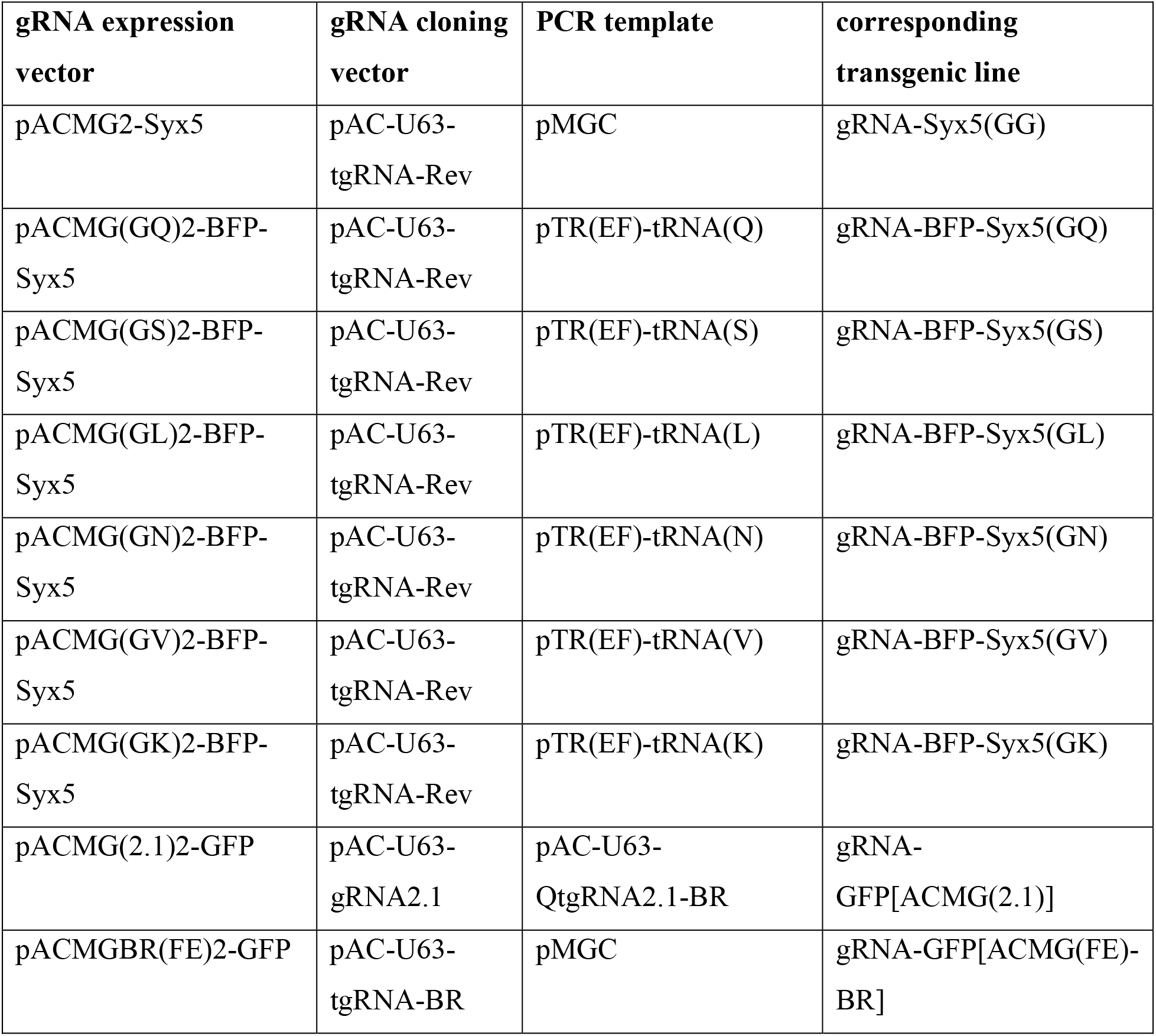
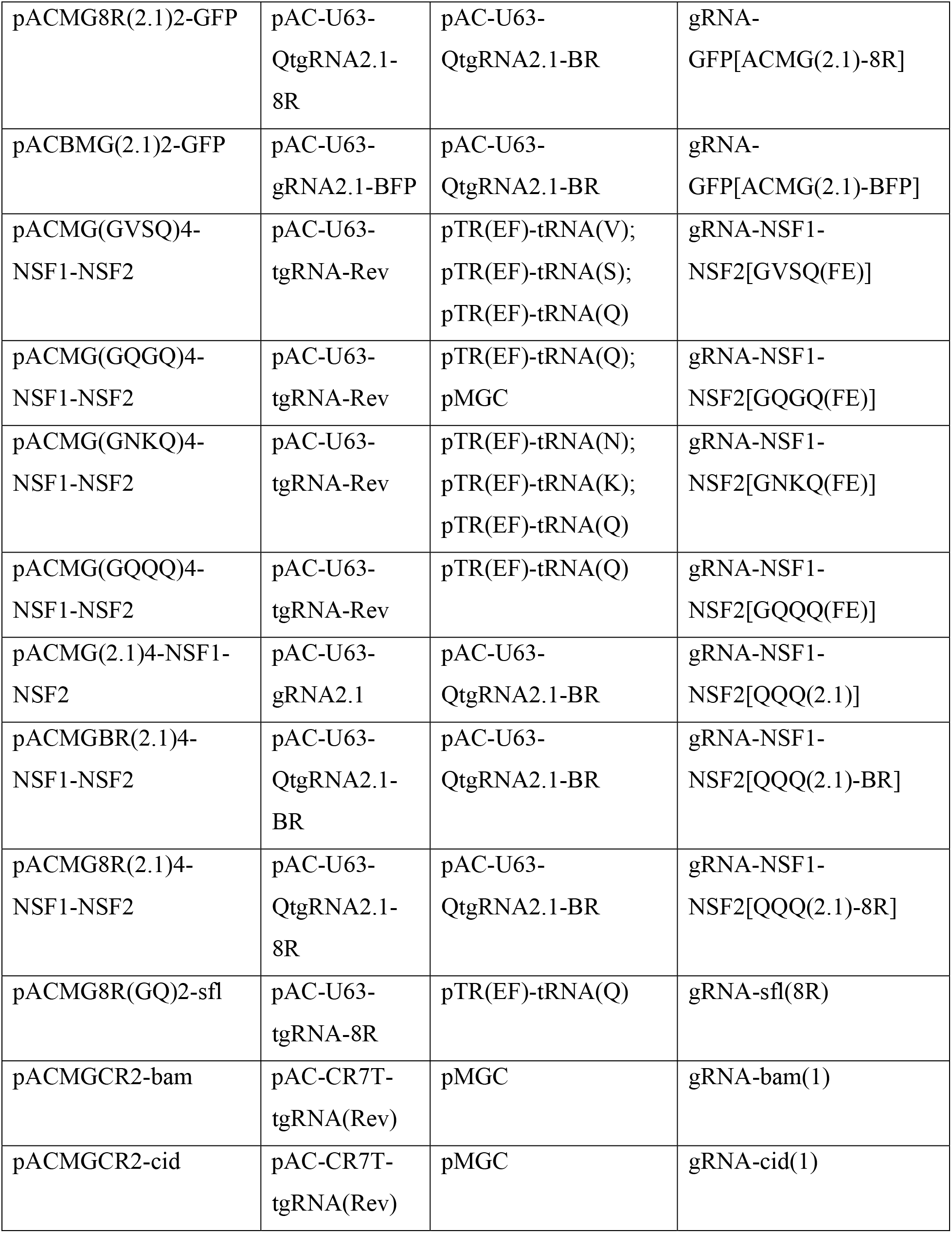
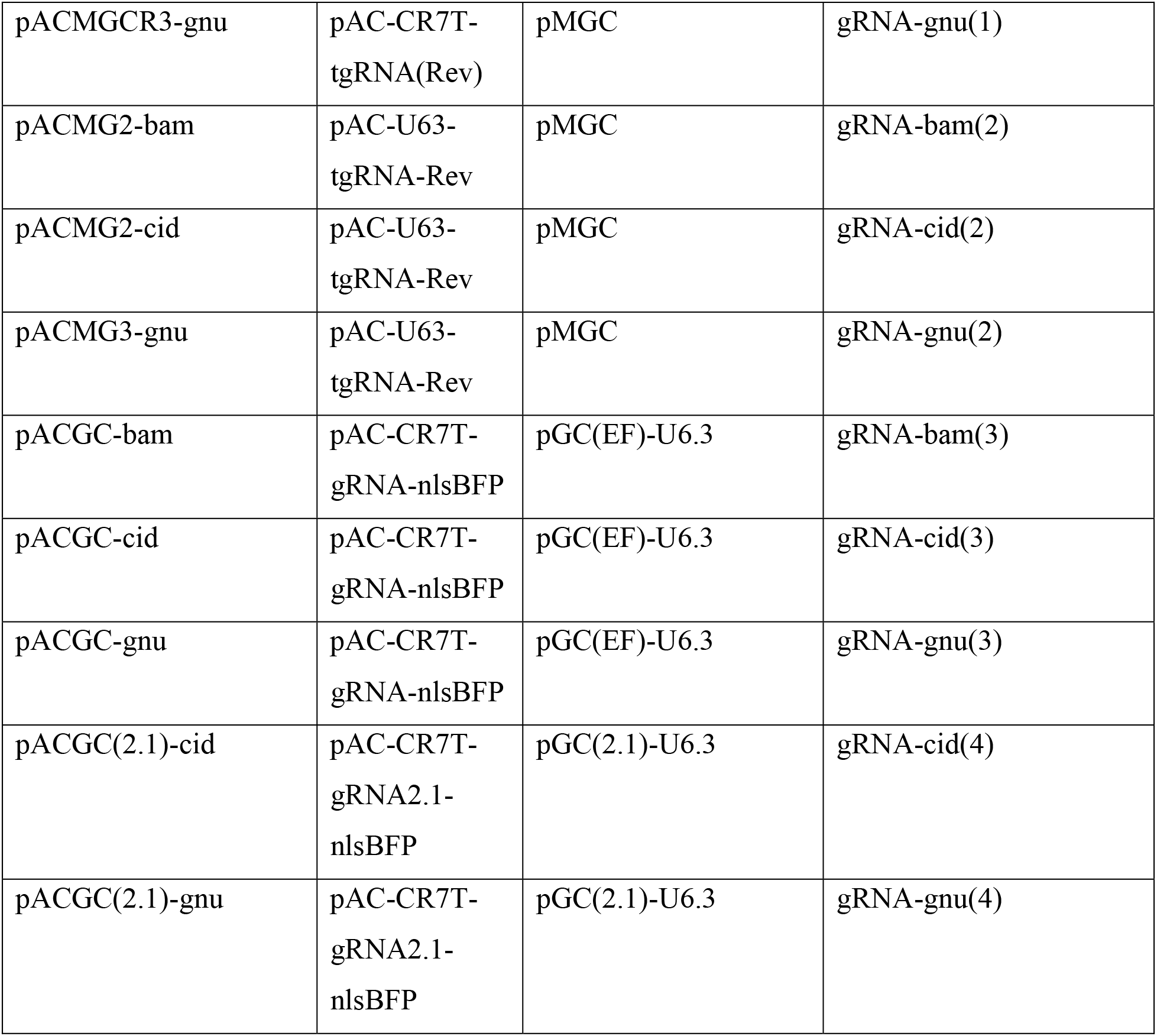
gRNA expression vectors.

**Table S6.**
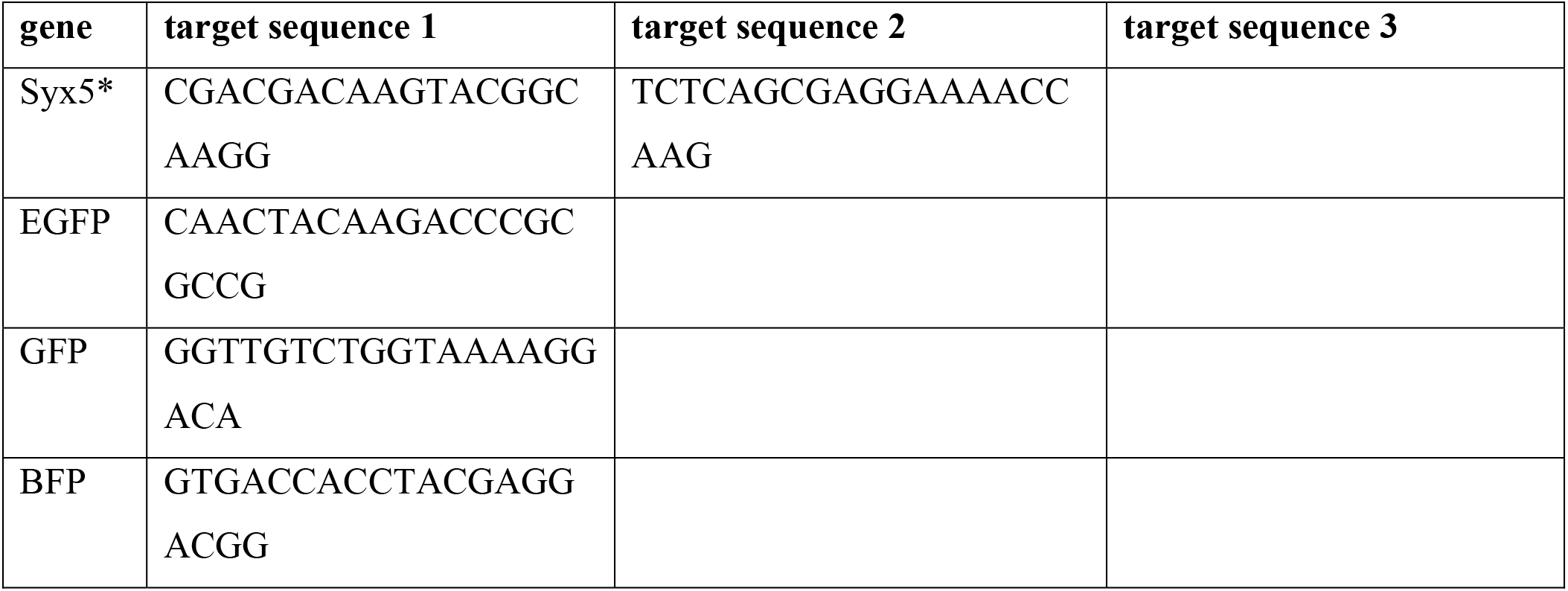
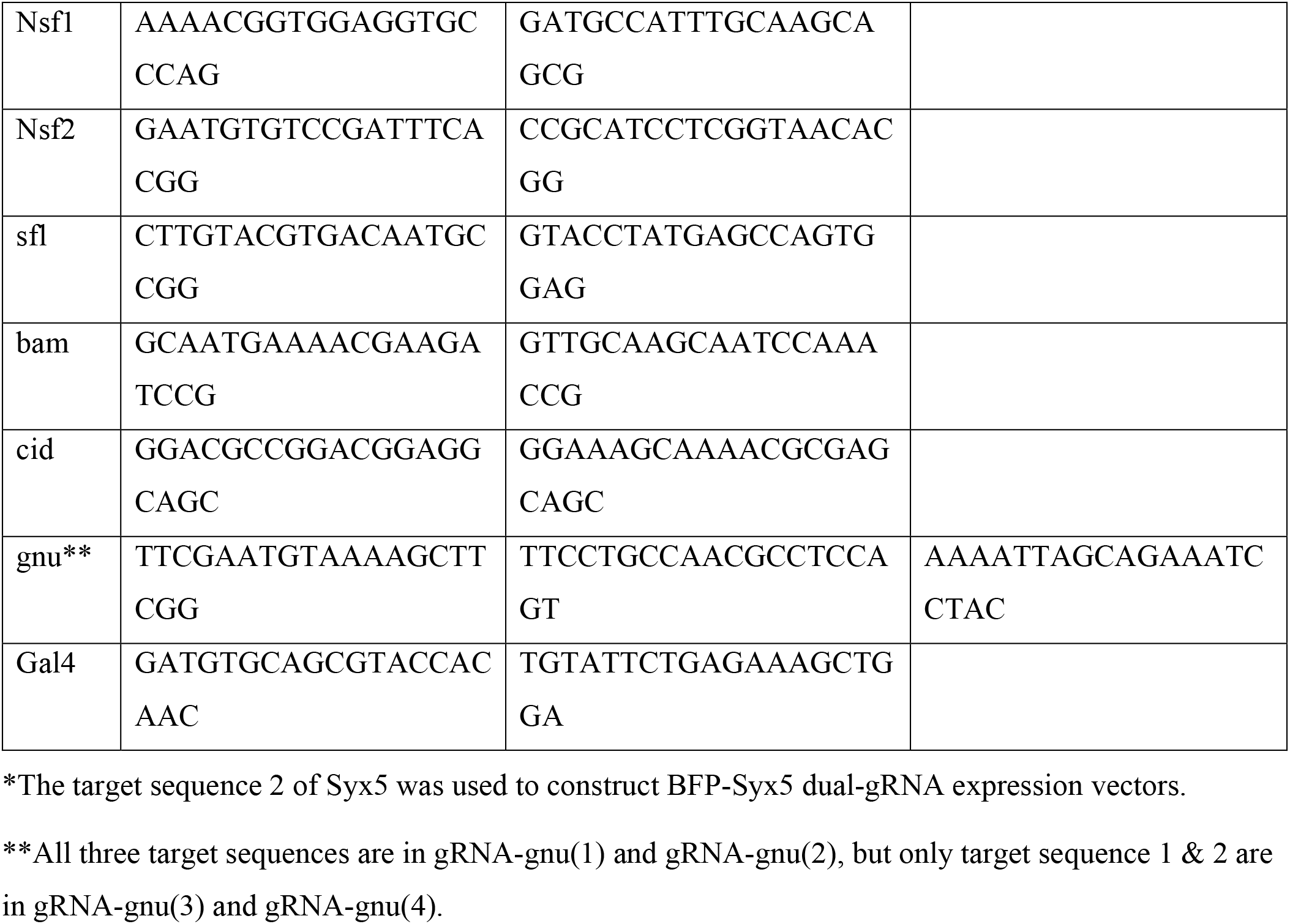
gRNA target sequences.

**Table S7.**
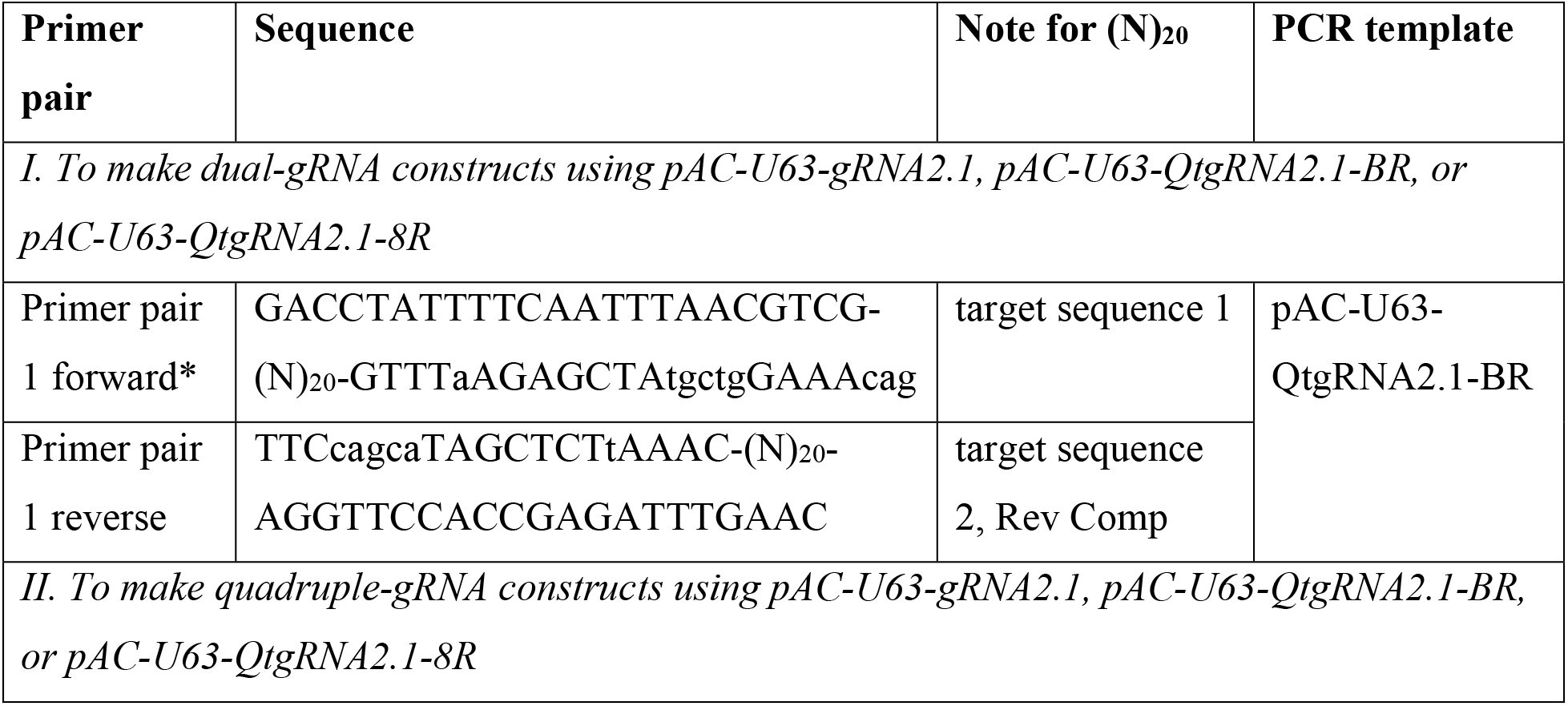
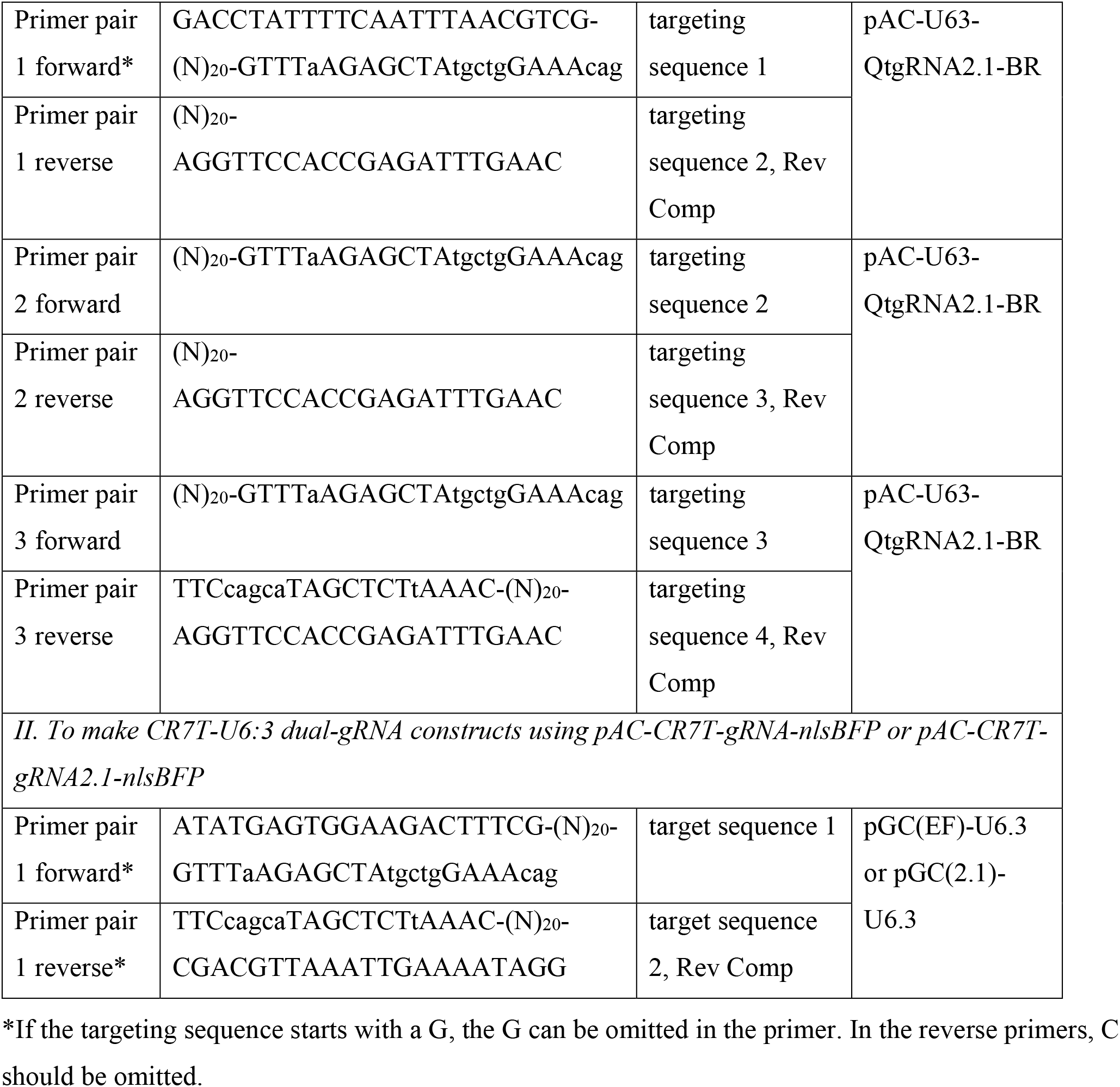
Primer designs for cloning gRNA expression constructs.

#### UAS-tdTom

The tdTomato coding sequence was PCR-amplified from pDEST-HemmarR2 (Addgene # 112813) and cloned into EcoRI/XbaI sites of pUA (Addgene 58372).

#### Gal4-to-Cas9 converter

Two gRNAs targeting Gal4 (34) were first cloned into pAC-CR7T-gRNA2.1-nlsBFP using pGC(2.1)-U6.3 as the PCR template, generating pACGC(2.1)-Gal4. Gal4 5’ and 3’ homology arms were PCR-amplified using pDEST-APIGH (Addgene # 112804) as the template. Cas9 coding sequence was PCR-amplified using pEDST-APIC-Cas9 (Addgene # 121657) as the template. These three PCR fragments were assembled together with a gBlock DNA fragment containing a hsp70 terminator sequence into PstI/NheI digested pACGC(2.1)-Gal4, resulting in the final construct pHACK(Gal4)-DONR(T2A-Cas9).

Injections were carried out by Rainbow Transgenic Flies (Camarillo, CA 93012 USA) to transform flies through φC31 integrase-mediated integration into attP docker sites. gRNA expression vectors were integrated into the *attP*^*VK00027*^ site. *UAS-tdTom* was integrated into the *attP*^*VK00033*^ site. The Gal4-to-Cas9 converter construct was integrated into *attP*^*40*^ and *attP*^*VK00027*^ sites. Transgenic insertions were validated by genomic PCR or sequencing.

All construct sequences are available upon request.

### Fertility assays

Virgin females of the indicated genotypes were aged on yeasted food and males were aged on non-yeasted food, both for 3 to 5 days. They were then mated with *Oregon-R-P2* (ORP2) (46) wildtype males or virgin females, respectively. Mating was observed and males were removed after a single mating had completed. Females were allowed to lay eggs in a mating vial for 24 hours and were transferred to a new vial. They were transferred 3 times before being discarded. Numbers of eggs and pupae were counted. Egg hatchability was calculated by the number of pupae divided by the number of eggs.

### Live Imaging

Live imaging was performed as previously described (28). Briefly, animals were reared at 25°C in density-controlled vials for between 96 and 120 hours (third to late-third instar). Larvae were mounted in glycerol and imaged using a Leica SP8 confocal microscope. The images of nuclear EGFP in the anterior half of third instar larvae were taken using a Nikon SMZ 18 stereomicroscope equipped with an Andor Zyla 3-Tap sCMOS Camera.

### Imaginal Disc/Brain imaging

Imaginal disc and larval brain dissections were performed as described previously (47). Briefly, wandering 3rd instar larvae were dissected in a small petri dish filled with cold PBS. The anterior half of the larva was inverted and the trachea and gut were removed. The sample was then transferred to 4% formaldehyde in PBS and fixed for 15 minutes at room temperature. After washing with PBS, the imaginal discs/brain were placed in SlowFade Diamond Antifade Mountant (Thermo Fisher Scientific) on a glass slide. A coverslip was lightly pressed on top. Discs were imaged using a using a Leica SP8 confocal microscope with a 40X oil objective while brains were imaged with a 20X oil objective.

### Immunohistochemistry

Following fixation, brains were rinsed and then washed twice at room temperature in PBS with 0.2% Triton-X100 (PBST) for 20 minutes each. Brains were then blocked in a solution of 5% normal donkey serum (NDS) in PBST for 1 hour. Brains were then incubated in the blocking solution with a mouse antibody mouse mAb NC82 (1:100 dilution, Developmental Studies Hybridoma Bank) for 2 hours at room temperature. Following incubation brains were then rinsed and washed in PBST 3 times for 20 minutes each. Brains were then incubated in a block solution containing a donkey anti-mouse secondary antibody conjugated with Cy5 (1:400 dilution, Jackson ImmunoResearch) and a rabbit anti-GFP antibody conjugated with Alexa Fluor 488 (1:500 dilution, Thermo Fisher Scientific) for 2 hours at room temperature. Brains were then rinsed and washed in PBST 3 times for 20 minutes each and stored at 4°C until mounting and imaging.

### Image Analysis and Quantification

Tracing and measurement of C4da dendrites was performed as described previously (Poe et al., 2017). Briefly, for tracing and measuring C4da dendrites in Fiji/ImageJ, images of dendrites (1,024 × 1,024 pixels) taken with a 20X objective were first processed by Gaussian Blur (Sigma: 0.8) and then Auto Local Threshold (Phansalkar method, radius: 50). Isolated particles below the size of 120 pixels were removed by the Particles4 plugin (http://www.mecourse.com/landinig/software/software.html). The dendrites were then converted to single-pixel-width skeletons using the Skeletonize (2D/3D) plugin and processed using Analyze Skeleton (2D/3D) plugin. The length of skeletons was calculated based on pixel distance.

Quantification of GFP(+) nuclei for the gRNA efficiency comparison was done in ImageJ/Fiji. ROIs were drawn from the anterior end to the segment A1 on images of the anterior half of third instar larvae taken using a Nikon SMZ fluorescent stereoscope at 5.2X magnification and 500ms exposure time. Images were processed using the Fiji subtract background function (rolling radius: 50), Gaussian Blur (Sigma: 1), and Auto Local Threshold (Phansalkar method, radius: 15). Particles of size above 35 pixels were isolated using the Analyze Particles FIJI function (circularity: 0.4-1.0). The total area of the selected particles was then divided by the total ROI area to give a percentage area of GFP coverage.

Quantification of GFP(+) and BFP(+) nuclei in epidermal cells for the co-CRISPR reliability assay was done in ImageJ/Fiji. BFP and GFP channels were separately processed to mask labeled nuclei using subtract background (rolling radius: 60), Gaussian Blur (sigma: 1) and auto local threshold (Phansalkar method, radius: 50). Particles of size above 80 pixels were counted as nuclei using the Analyze Particles. Images were then manually curated before quantified.

### Statistical Analysis

For dendrite length and EGFP(+) area analyses, when groups had equal variance and were normally distributed, a one-way analysis of variance (ANOVA) was performed followed by a Tukey’s honestly significant difference (HSD) test. When groups had unequal variances, Welch’s ANOVA was performed followed by post-hoc Welch’s t-tests with p-values adjusted using the Bonferroni method. Levene’s test was used to compare variances. For BFP-EGFP/EGFP ratio data and hatchability (hatched/not hatched) data, estimated marginal means (EMMs) contrasts were performed using a generalized linear mixed-effects models with binomial responses. Invariant groups, from the hatchability data, were compared using Fisher’s Exact Test. For egg laying data, EMMs contrasts were performed for each group based on a generalized linear model with a negative binomial response. p-values from all EMMs contrasts were adjusted using the Tukey method. All tests, correlation statistics (Pearson’s correlation coefficient), and linear regression models were generated using R.

---

**Figure S1:**
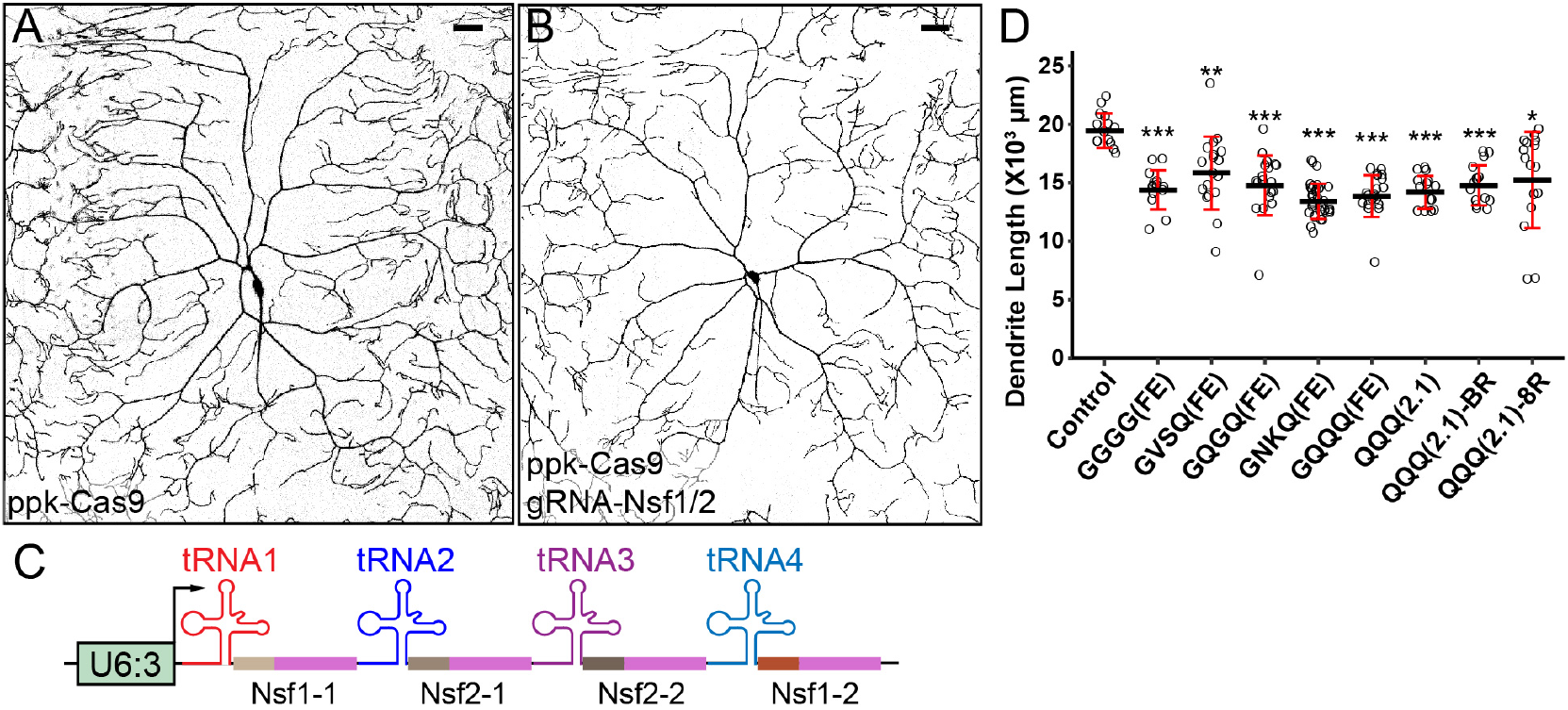
Performance of multi-gRNAs targeting *Nsf1* and *Nsf2*. (A and B) C4da neurons in *ppk-Cas9* control (A) and *ppk-Cas9 gRNA-NSF1-NSF2[GQQQ(FE)]* (B). (A) is the same as Figure 1A. Scale bars, 50 um. (C) General design of quadruple tRNA-gRNA constructs targeting *Nsf1* and *Nsf2*. (D) Quantification of C4da neuron total dendrite length using various gRNA-Nsf1-Nsf2 constructs. The significance level above each column indicates comparison with the control. *** p ≤ 0.001, **p≤ 0.01, *p ≤ 0.05; Welch’s ANOVA and Welch’s t-tests with Bonferroni correction. Each circle represents one neuron. n = number of neurons: Control (n = 14); GGGG(FE) (n = 14); GVSQ(FE) (n = 18); GQGQ(FE) (n = 19); GNKQ(FE) (n = 32); GQQQ(FE) (n = 20); QQQ(2.1) (n = 14); QQQ(2.1)-BR (n = 15); QQQ(2.1)-8R (n = 16). Black bar, mean; red bars, SD.

**Figure S2:**
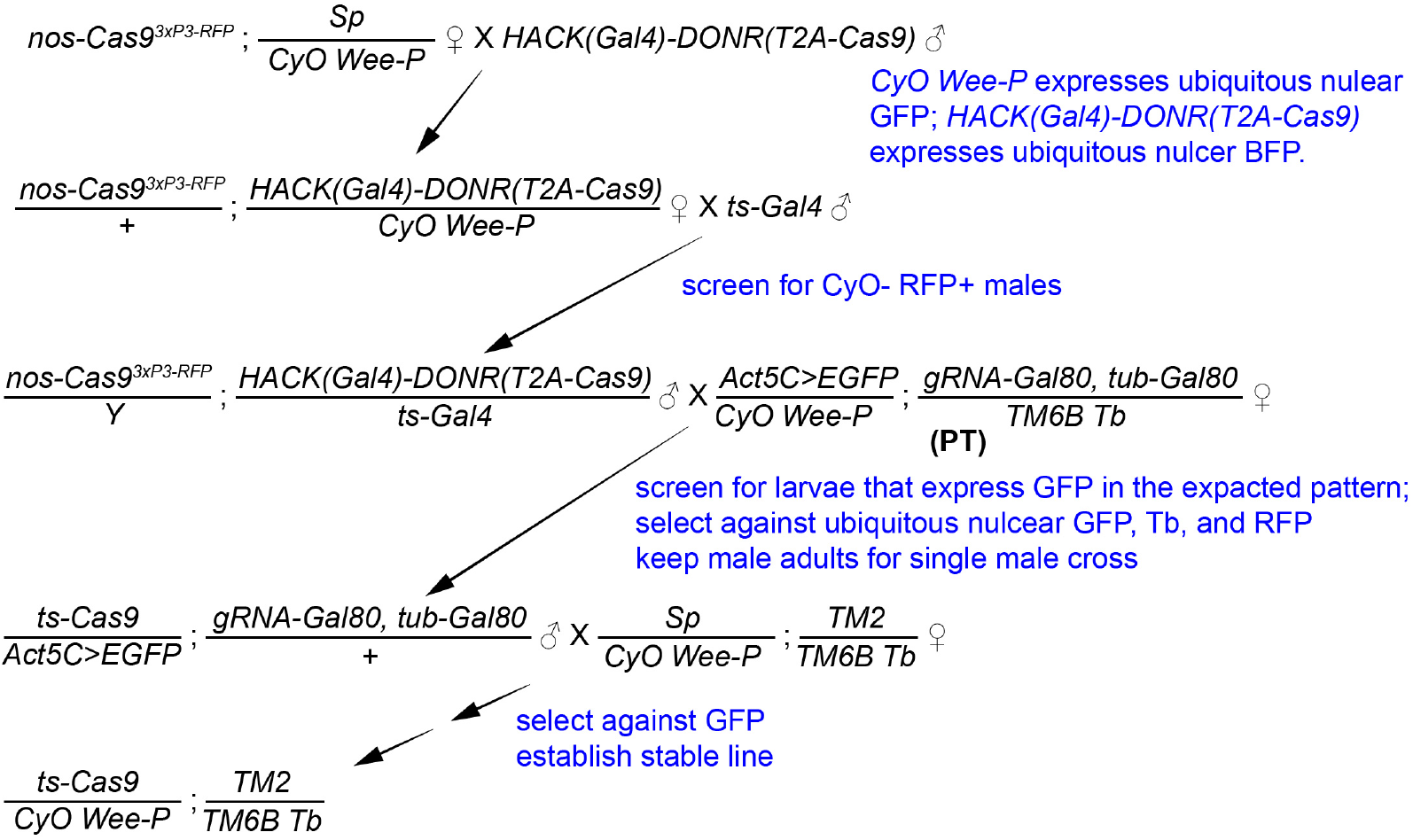
Steps for Gal4-to-Cas9 conversion. The steps illustrated are for conversion of a tissue-specific Gal4 line on the 2^nd^ chromosome.

## ACKNOWLEDGMENTS

We thank Bloomington *Drosophila* Stock Center (NIH P40OD018537) and KYOTO Stock Center for fly stocks; Cornell Statistical Consulting Unit (CSCU) for advice on statistics; Michael Goldberg for critical reading and suggestions on the manuscript. This work was supported by a Cornell start-up fund and NIH grants (R01NS099125 and R21OD023824) awarded to C.H., R21-HD088744 awarded to M.F.W‥

## AUTHOR CONTRIBUTIONS

G.T.K., Q.H., Y.X., M.F.W., B.W. and C.H. designed research; G.T.K., Q.H., Y.X., Z.Z., and S.A. performed research; Q.H. and B.W. contributed new reagents/analytic tools; G.T.K., Q.H., Y.X., M.F.W., B.W. and C.H. analyzed data; G.T.K., Q.H., Y.X. and C.H. wrote the manuscript; all authors edited the manuscript; M.F.W. and C.H. acquired funding.

